# Vimentin enables directional cell migration by coordinating focal adhesion organization and dynamics

**DOI:** 10.1101/2022.10.02.510295

**Authors:** Arun P. Venu, Mayank Kumar Modi, Elena Tcarenkova, Giulia Sultana, Jenny Pessa, Peiru Yang, Ujjwal Aryal, Leila Rato, Joakim Edman, Yaming Jiu, Guillaume Jacquemet, Alexander Minin, Fang Cheng, John E. Eriksson

## Abstract

Persistent cell migration requires focal adhesions to assemble and disassemble locally while maintaining global front-rear alignment. The mechanism that enforces this long-range spatial coherence remains unresolved. Here we identify the intermediate filament protein vimentin as a cell-scale organizer that stabilizes focal adhesion alignment during directed fibroblast migration. Using quantitative live-cell imaging, we show that vimentin-deficient fibroblasts lose directional persistence and a complete collapse of global focal adhesion alignment. Quantitative analysis reveals that vimentin stabilizes focal adhesion alignment by constraining angular fluctuations and preserving the periodic bias of adhesion birth across the adhesion field. Loss of vimentin results in smaller, rapidly turning-over adhesions with disrupted orientation. Trajectory analysis reveals a mechanically anchored adhesion state selectively associated with vimentin recruitment, distinguishing mechanical stabilization from biochemical maturation. Super-resolution and iPALM imaging further show that vimentin integrates within the focal adhesion nanoarchitecture near the force-transduction layer. Together, our findings establish that vimentin intermediate filaments impose spatial coherence on adhesion dynamics, converting locally stochastic adhesion assembly, turnover, and disassembly into globally coordinated adhesions and persistent directional migration.

## Introduction

Cell migration is essential to a range of key physiological processes, such as immune response, wound healing, cancer metastasis, and embryonic development (Chernoivanenko et al.,2013; Conway & Jacquemet, 2019). Cell migration is enabled through highly dynamic and concerted interactions between cytoskeletal and adhesion proteins, the joint maneuvering of which regulates adhesive properties of cells, determine cell polarity, and the speed and direction of cells. Directional cell migration is maintained by the coordinated interplay between different cytoskeletal systems and their interactions with focal adhesions (FAs), which enable a spatiotemporally sustained movement of the cellular projections at the leading edge (Seetharaman & Etienne-Manneville, 2020).

The intermediate filament (IF) protein vimentin has been shown to be involved in cell migration in a number of instances that relate to wound healing and cancer progression (Battaglia et al., 2018; Cheng et al., 2016, 2017; Coelho-Rato el al. 2024; Satelli & Li, 2011). Vimentin is expressed in fibroblasts, endothelial cells, lymphocytes, and several specialized cells, all of which are characterized by active adhesion-dependent migratory functions and all of which arise from the migratory cells of developing embryos. Vimentin has also been established as a hallmark of epithelial to mesenchymal transition in several physiological processes (Cheng et al., 2016; Coelho-Rato et al. 2024; Ivaska et al., 2011; Virtakoivu et al., 2015).

While vimentin knockout experiments initially revealed a mild phenotype in Vim^−/−^ mice (Colucci-Guyon et al., 1994), subsequent studies have shown that these mice exhibit a broad spectrum of distinct phenotypes, involving a wide range of deficiencies related to maintenance of stemness, differentiation, proliferation, mammary gland development, angiogenesis, vascular stiffness, steroidogenesis, glia development, myelination of peripheral nerves, adhesion, migration and invasion (Cheng & Eriksson, 2017; Ridge et al. 2022, Lusk & Eriksson, 2024). The mice have a prominent defect in wound healing (Cheng et al., 2016; Eckes et al., 2000; reviewed in Cheng and Eriksson 2017; Coelho-Rato et al. 2024; Ridge et al. 2022), which has been shown to be due to strongly impaired fibroblast growth and migration at the wound edge, as well as inhibited TGF-β signaling, and epithelial to mesenchymal transition (Cheng et al., 2016). Furthermore, Vim^−/−^ fibroblasts have exhibited delayed migration towards the wound edge (Cheng et al., 2016; Eckes et al., 2000). Leukocytes, astrocytes, and different types of cancer cells also show improper motility when vimentin is absent (Satelli & Li, 2011).

Directional orientation is important for motile cells to reach a particular destination inside the organism in response to signaling cues (Ridley et al., 2003). A key determinant of directional migration is the establishment and maintenance of cell polarity (Etienne-Manneville et al., 2013; Zhang et al., 2014). Several studies have shown that intermediate filaments play key roles in regulating cell migration, cell polarity, and cell-ECM interactions in fibroblasts (Ivaska et al., 2007; Jiu et al., 2015; Jiu et al., 2017), largely through crosstalk with actin cytoskeleton dynamics and Rho-family signaling pathways (Jiu et al., 2015; Jiu et al., 2017; Seetharaman & Etienne-Manneville, 2020). In collectively migrating astrocytes expressing vimentin, GFAP, and nestin, disruption of the intermediate filament network impairs long-range force transmission and directional coherence (De Pascalis et al., 2018). (De Pascalis et al., 2018). However, how these cytoskeletal and signaling functions are integrated at the level of adhesion dynamics to generate persistent directional migration remains unclear.

Beyond these established cytoskeletal interactions, recent work has further underscored the importance of vimentin in coordinating collective cell migration and tissue-scale repair. Strikingly, a recent study demonstrates that vimentin is essential for long-range coordination and force redistribution during collective epithelial migration, highlighting the timeliness and importance of understanding how vimentin confers directional order at the tissue level (Peneti et al., 2026). Furthermore, the expression of vimentin has recently been shown to be a critical driver of 3D multicellular invasion by enhancing cell-matrix interactions and actively localizing traction forces (Rodriguez et al., 2026). Together, these findings raise a central unresolved question: how does vimentin, at the molecular and subcellular level, interface with adhesion-based force transmission to impose persistent, directional migration.

Given the previously indicated multifactorial roles of vimentin in migrating cells and that IFs in general have been implicated as important determinants of cell migration, we wanted to specifically address the role of vimentin in fibroblasts directionality and we sought to define the mechanistic principles by which vimentin regulates directional persistence in fibroblasts. Fibroblasts have a straightforward IF composition, with vimentin as the only type-III IF protein. We observed a pronounced loss of directionality in mouse and rat embryonic fibroblasts. As FA organization and control are key for directed migration, we wanted to specifically address how vimentin regulates FAs during directional cell migration. Although vimentin has been shown to associate with FAs and in some cell types to interact with integrins (Burgstaller et al., 2010; Ivaska et al. 2005; Strouhalova et al., 2020), the mechanistic links between vimentin cytoskeleton, cell adhesion dynamics, and the cellular forces acting on the ECM remain poorly understood. Here, we show that vimentin regulates directional cell migration by controlling the dynamics, orientation, and coordinated organization of FAs across the cell.

## Results

### Vimentin regulates directional cell migration

To examine the effects of vimentin on directional migration required for wound healing in detail, we performed scratch wound assays using WT and Vim KO mouse embryonic fibroblasts MEFs to replicate the wound healing process at the cellular level in vitro and to induce polarization of cells that become stimulated to migrate ( Video 1A-left panel). The cells were allowed to migrate for 24 hours, and single cells were then manually tracked. Only cells migrating perpendicular to the wound edge from left to right were included in the analysis, and their trajectories were used to calculate migration directionality and directional persistence. WT MEFs exhibited robust, persistent migration oriented along the wound axis, with most cells displaying straight trajectories and clear front-rear polarity (Fig. 1A; Video 1A, left panel). In contrast, vimentin-deficient (Vim^−/−^) MEFs exhibited a complete loss of directional persistence, (Fig. 1B; Video 1A, right panel), with migration paths appearing disorganized and frequently changing direction. This loss of directional persistence was also observed in rat embryonic fibroblasts (REFs) lacking vimentin (Fig. S2B, Video 1B, right panel), demonstrating that the requirement for vimentin in directional migration is conserved across species.

**Figure 1.**
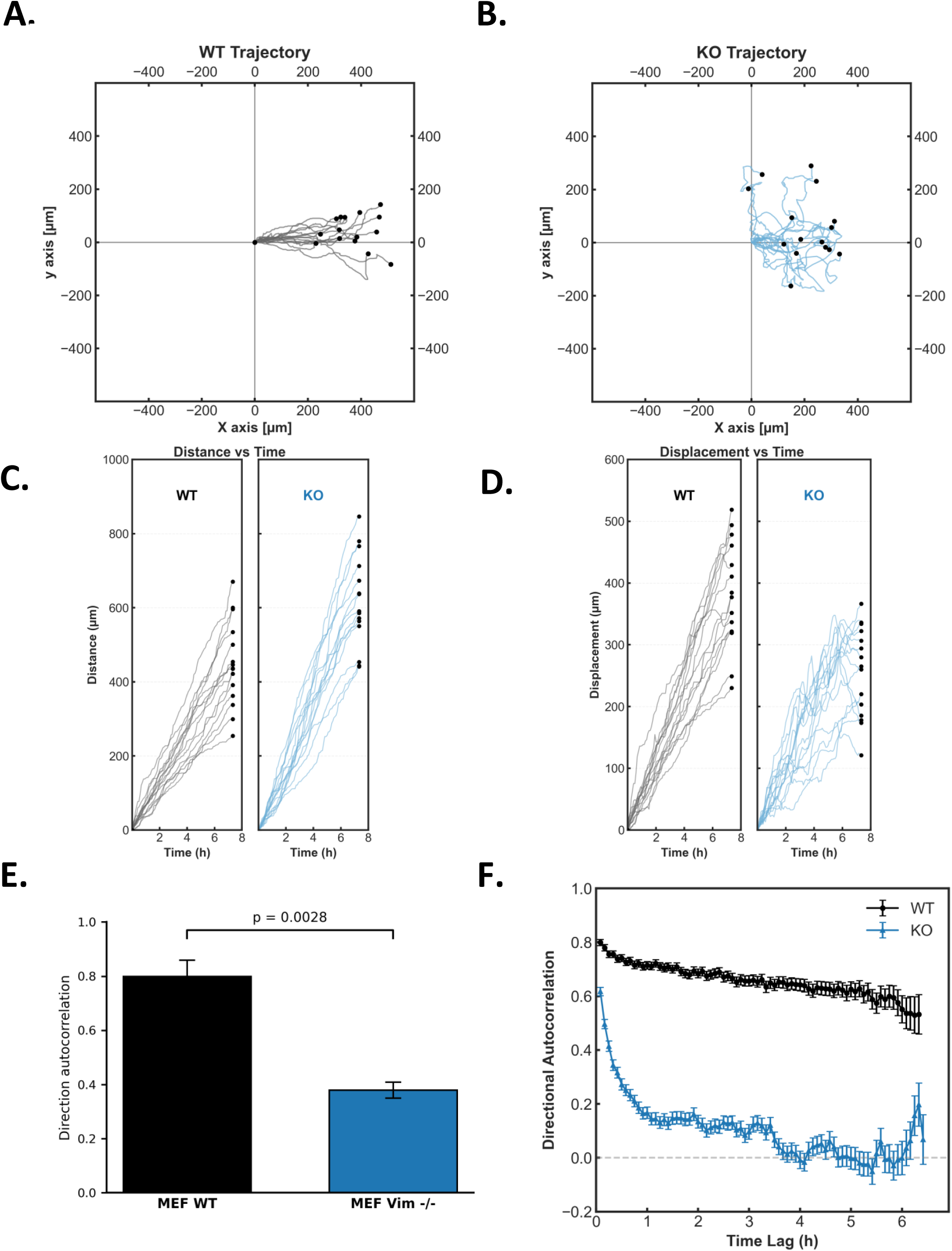
Directional migration is lost in vimentin-deficient fibroblasts. (A) Representative single-cell migration trajectories of wild-type (WT) mouse embryonic fibroblasts (MEFs) during scratch wound migration, aligned to a common origin and wound axis. (B) Corresponding trajectories of vimentin-deficient (Vim^−/−^) MEFs show disorganized and non-persistent migration. (C) Total migration distance as a function of time for individual WT and Vim^−/−^ MEFs. (D) Net displacement over time for WT and Vim^−/−^ MEFs. (E) Directionality ratio analysis reveals significantly reduced directional persistence in Vim^−/−^ MEFs compared to WT cells. (F) Time-lag–dependent directional autocorrelation demonstrates rapid loss of directional memory in Vim^−/−^ MEFs. Data were obtained from 15 cells per condition across three independent experiments.

Intriguingly, the loss of directional migration in Vim^−/−^ fibroblasts was not related to reduced motility. Instead, while Vim^−/−^ MEFs had dramatically reduced directionality score compared to WT (Fig. 1E), Vim^−/−^ MEFs migrated at increased speeds (Fig. S1B) and covered longer total path lengths than WT cells (Fig. 1 C, Fig. S1C). While moving at higher speed, they displayed significantly reduced net displacement from the origin (Fig. 1 D). Thus, despite moving rapidly, Vim^−/−^ fibroblasts followed highly erratic, non-persistent trajectories, resulting in a marked reduction in their displacement, which was in stark contrast to the sustained and directed migration of WT cells. These data demonstrate that loss of vimentin specifically disrupts directional control and persistence, uncoupling migratory speed from directional migration capacity.

To quantitatively assess directional migration efficiency, we calculated directionality as the ratio of net displacement to total migration distance. WT MEFs displayed significantly higher directionality than Vim^−/−^ MEFs (Fig. 1E; p = 0.0028), reflecting straighter and more persistent migration paths. In contrast, Vim^−/−^ MEFs exhibited erratic, meandering trajectories with reduced directional efficiency.

To quantify directional persistence over time, we computed direction autocorrelation of migration vectors. WT fibroblasts maintained high autocorrelation values over extended time lags, indicating stable directional persistence (Fig. 1F). In contrast, Vim^−/−^ MEFs showed a rapid decay in autocorrelation, reflecting frequent reorientation and loss of stable migration direction. These complementary analyses demonstrate that vimentin is critical for both the efficiency and the temporal stability of directional cell migration, supporting its essential role in maintaining front-rear polarity and coordinated movement.

To determine whether the loss of directionality observed in wound healing assays is specific to polarized migration or reflects a more general defect in motility, we next examined random migration in sparsely plated fibroblast cultures. Under these conditions, WT MEFs and REFs exhibited extended trajectories with sustained directional persistence (Fig. S1D; Fig. S2C). In contrast, Vim^−/−^ MEFs and REFs displayed short, erratic trajectories with frequent turning and limited net displacement (Fig. S1E; Fig. S2D), demonstrating that directional defects occur independently of imposed polarization cues.

To investigate whether matrix-guided cues contribute to the observed directional deficiency phenotype, we seeded WT and vimentin-deficient (Vim^−/−^) MEFs on cell-derived matrix (CDM) produced by WT fibroblasts. On this biologically relevant substrate, WT MEFs exhibited highly directed migration, with long, outwardly oriented trajectories that aligned with the underlying matrix fibers (Fig. S1F, Video 2, left-hand panel). In contrast, Vim^−/−^ MEFs displayed disorganized, short, and erratic paths (Fig. S1G, Video 2, right-hand panel), lacking the alignment or persistence observed in WT cells and also lacking the alignment with CDM. Strikingly, the movement of Vim^−/−^ cells appeared to disturb and disorganize the CDM itself Video 2, suggesting failure to engage with matrix guidance cues.

Together, these results demonstrate that vimentin is essential for maintaining intrinsic directional migration capacity of fibroblasts across multiple migratory contexts, including polarized wound closure, random migration, and matrix-guided migration. Vimentin, thus, supports directional persistence, front-rear polarity, and adhesion dynamics during migration, and its loss results in defective matrix-guided migration accompanied by disruption of cell-derived matrix organization.

### Vimentin is required for focal adhesion alignment during directed migration

FAs are critical for establishing front-rear polarity and guiding directional migration by serving as mechanical anchor points and signaling hubs that align cytoskeletal dynamics with the direction of movement. Given this essential role of FAs and the dramatic loss of directional migration observed in vimentin-deficient cells, we hypothesized that the absence of vimentin may also disrupt the spatial orientation and dynamic behavior of FAs during migration.

To investigate this, WT and Vim^−/−^ MEFs expressing Emerald-Paxillin were imaged by live-cell TIRF microscopy during scratch wound migration (Video 3). Over 2000 individual adhesions were tracked and analyzed from time-lapse Videos acquired at 20-second intervals over 1 hour (Video 3). Leader cells at the wound edge were selected for analysis, and time-lapse sequences were analyzed using the Focal Adhesion Analysis Server (FAAS) (Berginski & Gomez, 2013).

To assess whether vimentin influences FA alignment, FA orientation over time was visualized by superimposing all time-lapse frames from individual migrating cells (Video 3). As shown in Fig. 2A (left panel), WT MEFs exhibited a highly coordinated and aligned progression of FA formation, with adhesions forming and maturing along consistent spatial axes, from initial (blue) to final (red) time points. In stark contrast, Vim^−/−^ MEFs displayed disorganized and randomly oriented FA dynamics, as evidenced by the lack of uniformity in FA development trajectories over time (Fig. 2A, right panel; Video 3). This disordered pattern is indicative of a failure in global FA alignment and directionally consistent FA development.

**Figure 2.**
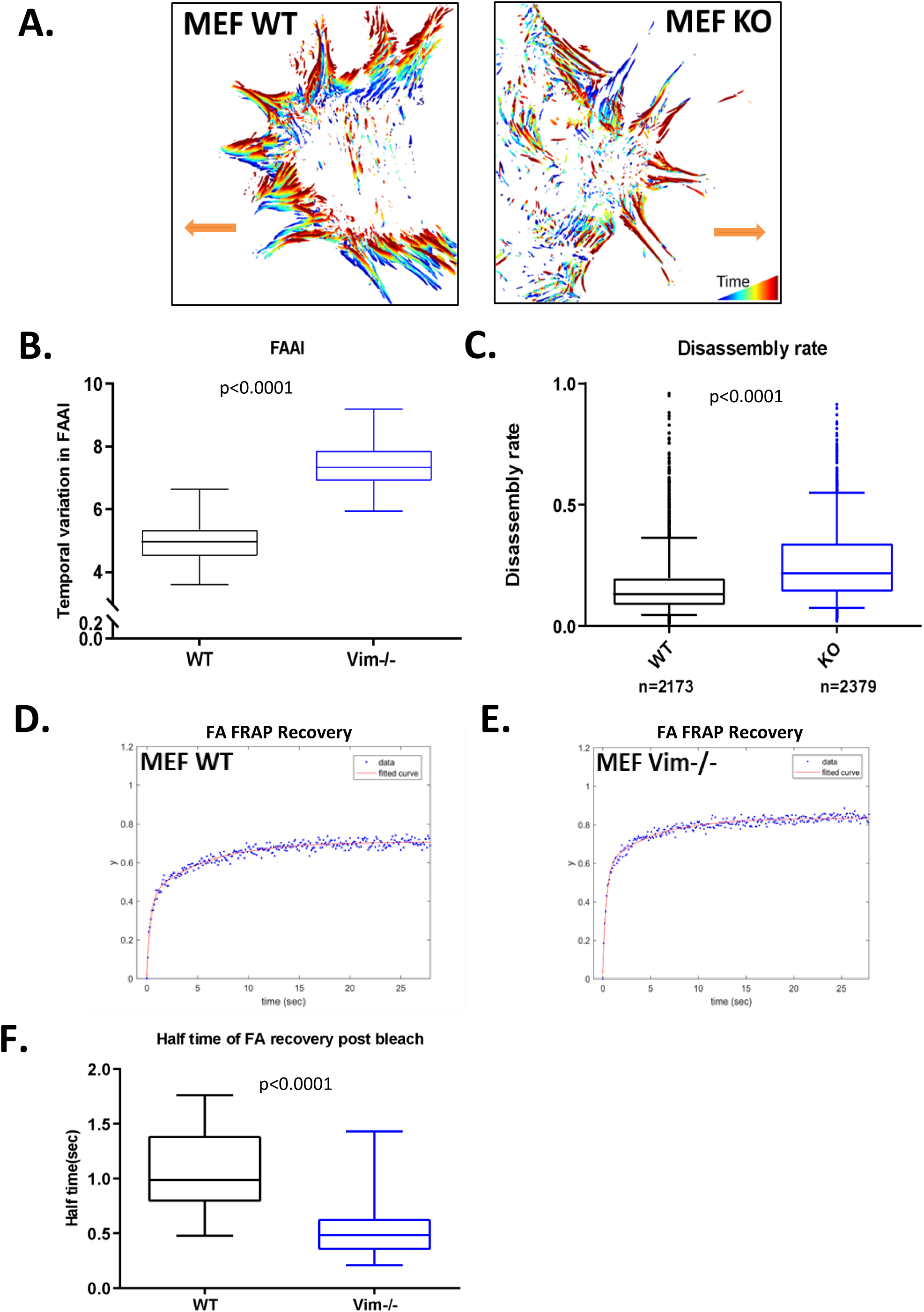
Vimentin deficiency causes disorganized and unstable focal adhesions during directional migration. (A) Superimposed time-lapse frames of focal adhesions (FAs) in WT and Vim^−/−^ MEFs expressing Emerald-paxillin during wound-directed migration. Color progression from blue to red represents temporal evolution of FA dynamics. (B) Temporal variation of the focal adhesion alignment index (FAAI) shows increased variability in Vim^−/−^ MEFs. (C) Distribution of FA disassembly rates reveals significantly increased disassembly in Vim^−/−^ cells (n > 2000 adhesions; 30 cells; three independent experiments). (D–E) Mean fluorescence recovery after photobleaching (FRAP) curves of individual FAs in WT (D) and Vim^−/−^ (E) MEFs. (F) Quantification of FRAP recovery half-times demonstrates increased paxillin turnover in Vim^−/−^ FAs. Imaging was performed using TIRF microscopy at 20-s intervals.

When analyzing the population-level variation of the FA alignment index (FAAI), calculated as 90° minus the standard deviation (SD) of FA angles, we found that Vim^−/−^ MEFs (Fig. 2B) exhibited significantly greater variation in FAAI compared to WT MEFs (Fig. 2B), consistent with the disorganized FA alignment observed visually in Fig. 2A.

We also found that WT MEFs displayed a relatively stable FAAI over time, with only minor fluctuations around a consistent mean, indicating well-coordinated and persistent FA alignment during migration (Fig. S3A). In sharp contrast, Vim^−/−^ MEFs exhibited pronounced temporal variability in FAAI, reflecting unstable and poorly coordinated FA orientation (Fig. S3B). Notably, FAAI fluctuations in Vim^−/−^ cells were approximately twofold greater than in WT cells, indicating a profound loss of temporal control over FA alignment in the absence of vimentin.

The disruption of FA coordination in Vim^−/−^ cells is functionally linked to the loss of directional persistence during migration, consistent with a role for vimentin in stabilizing FA orientation over time. Collectively, these findings indicate that vimentin is required to maintain the directional orientation and temporal stability of FA dynamics during directed migration.

This conclusion is further supported by analysis of FA birth angles during FA formation (Fig. S3C). In WT MEFs, newly formed FAs exhibited a clear periodicity in their birth angles, indicating preferential emergence at defined orientations relative to the cell’s migratory axis. In contrast, Vim^−/−^ MEFs lacked this periodic organization and displayed a near-uniform angular distribution of FA birth events. The loss of angular bias in FA formation suggests that vimentin is required to impose spatial directionality on nascent adhesions. Consequently, the reduced FAAI variability observed in WT cells likely arises from vimentin-dependent coordination of FA birth orientation. Together, these results demonstrate that vimentin is essential for maintaining global FA alignment and angular periodicity during migration.

In summary, our results establish that vimentin is required for global FA alignment and stabilization of FA orientation dynamics during directed migration. In its absence, FAs lose coordinated alignment, exhibit pronounced temporal instability in FAAI, and fail to maintain the biased, periodic birth-angle organization characteristic of polarized migration. These defects disrupt the spatial coordination of adhesion remodeling and impair the alignment of traction forces, ultimately resulting in loss of persistent directional migration. Together, our findings identify vimentin as a previously unrecognized regulator of FA orientation dynamics over time that is essential for maintaining directional migration and front-rear polarity.

### Vimentin stabilizes focal adhesions by limiting turnover and disassembly dynamics

Building on the observation that FA orientation and coordination are disrupted in vimentin-deficient cells, we next investigated whether vimentin also modulates the stability and turnover dynamics of individual FAs during migration.

To this end, we used the above-mentioned analysis of WT and Vim^−/−^ MEFs expressing Emerald-Paxillin imaged by live-cell TIRF microscopy during scratch wound migration (Video 3), including over 2000 individual adhesions (Video 3). FA assembly and disassembly rates were quantified from changes in paxillin fluorescence intensity over time using custom image-analysis pipelines (n = 30 cells, 3 independent experiments). FA assembly and disassembly rates were calculated based on the slope of fluorescence intensity increase or decrease over time, corresponding to the growth or shrinkage phase of each individual FA.

While the assembly rate of FAs was comparable between WT and Vim^−/−^ MEFs (data not shown), the disassembly rate was significantly increased in Vim^−/−^ cells (Fig. 2C), indicating accelerated FA turnover in the absence of vimentin. This suggests that although adhesions can initially form at a normal rate, their subsequent stability and lifetime are compromised when the vimentin cytoskeleton is absent. To directly assess adhesion stability, we performed fluorescence recovery after photobleaching (FRAP) experiments on individual FAs in MEF WT and Vim^−/−^ cells expressing Emerald-Paxillin. Vim^−/−^ MEFs exhibited significantly faster fluorescence recovery than WT cells (Fig. 2D, E), indicating increased molecular turnover within the FAs. Quantification of recovery half-times revealed that the paxillin in the Vim^−/−^ FAs recovered nearly twice as fast as in WT adhesions (Fig. 2F). This reflects increased paxillin turnover in the Vim^−/−^ FAs, which is consistent with the assumption of reduced FA stability.

Taken together, these findings demonstrate that vimentin is essential for stabilizing FAs by limiting their disassembly and molecular turnover, consistent with the established role of vimentin as a dynamic, phosphorylation-regulated modulator of cellular plasticity (Eriksson et al., 1992; Eriksson et al., 2004; Hyder et al., 2011; Coelho-Rato et al., 2024). The accelerated turnover observed in Vim^−/−^ cells disrupts the FA persistence and coordination necessary for directional migration. The loss of sustained FA behavior across the migrating cell likely underlies the observed loss of directional persistence. The observed increased FA dynamism is likely to contribute to the elevated migration speed observed in Vim^−/−^ cells (see Fig. 1, Fig. S1B), enabling faster cells that have completely lost their direction of movement.

### Vimentin deficiency reduces FA size but not the number of FAs

Directional migration requires the coordinated assembly, maturation, and disassembly of FAs. To explore how vimentin contributes to this process, we analyzed the structural organization of FAs in migrating WT and Vim^−/−^ fibroblasts (Fig. 3A). Cells were fixed 4 hours after wounding in a scratch assay and immunostained for paxillin. Confocal imaging revealed that while the number of FAs per cell was similar in both genotypes (Fig. 3B), the average FA area was significantly smaller in Vim^−/−^ cells (Fig. 3C), indicating defective FA maturation.

**Figure 3.**
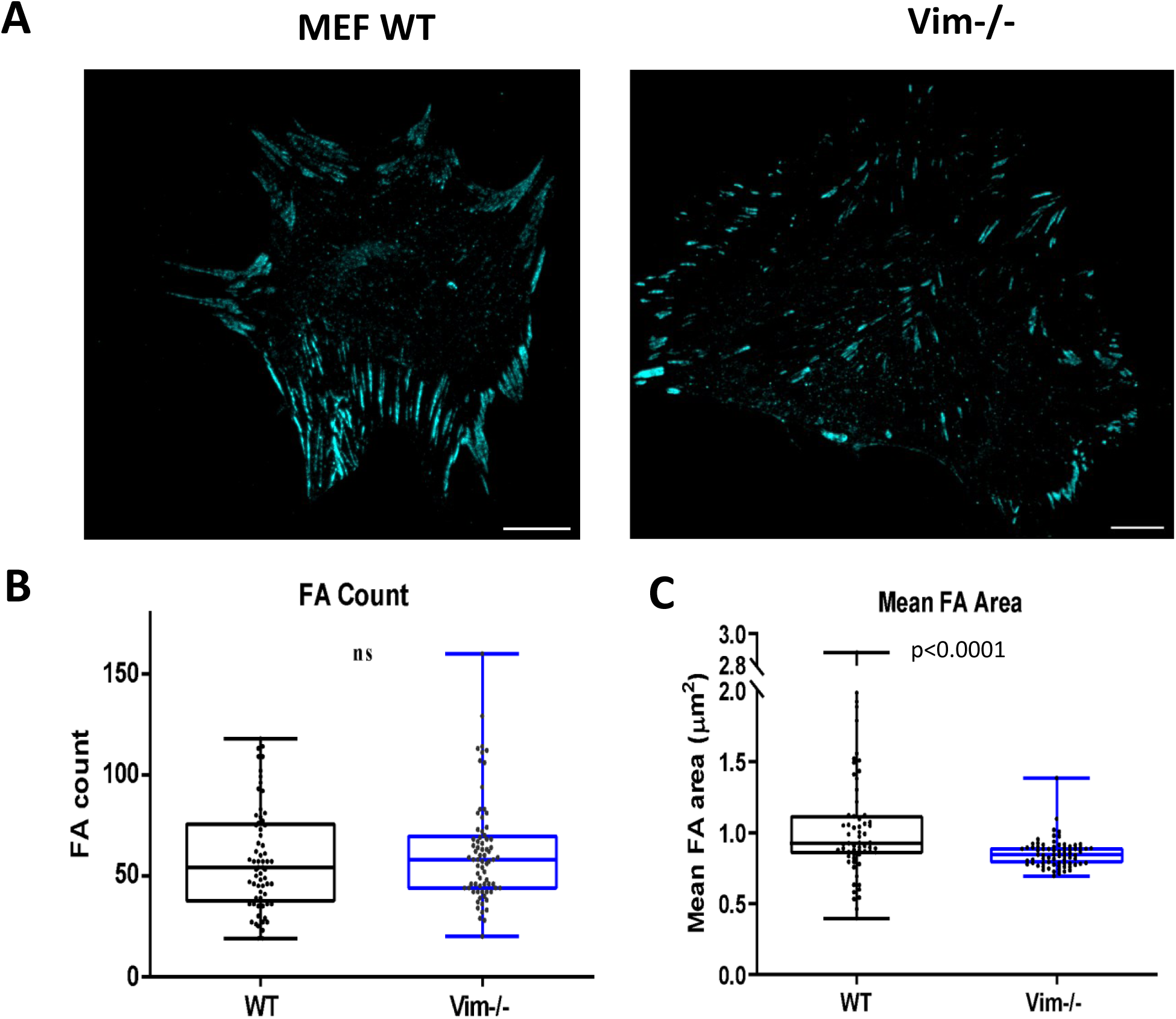
Vimentin deficiency reduces focal adhesion area but not focal adhesion number. (A) Representative spinning-disk confocal images of paxillin-stained FAs in WT and Vim^−/−^ MEFs at the wound edge (maximum-intensity projections). (B) Quantification of FA number per cell shows no significant difference between genotypes. (C) Mean FA area is significantly reduced in Vim^−/−^ MEFs, indicating impaired FA maturation. Scale bar, 10 µm.

These findings are consistent with the increased disassembly rate observed in Vim^−/−^ cells (Fig. 2C) and suggest that vimentin stabilizes FAs not only temporally but also structurally by promoting FA growth and retention. The reduced FA size may reflect premature disassembly or impaired maturation, both of which compromise adhesion strength and directional traction during migration.

Taken together, these results demonstrate that the loss of vimentin leads to smaller and less stable FA. This suggests that vimentin supports FA maturation and stability by regulating disassembly dynamics. A faster FA turnover rate in KO can lead to a loss of accumulation, leading to lower FA size.

### Vimentin interacts with maturing and retracting adhesions

To investigate whether the observed disruption in FA alignment in vimentin-deficient cells could be explained by direct interactions between vimentin and FA components, we analyzed the spatiotemporal relationship between vimentin filaments and FAs during migration.

As FAK is such a key regulator of FAs, we first studied the reciprocal spatial organization and dynamics between vimentin and FAK. To this end, live-cell imaging of Vim^−/−^ MEFs reconstituted with with mCherry-tagged vimentin and co-transfected with GFP-tagged FAK revealed dynamic interactions between vimentin and FAs. TIRF imaging demonstrated that short and long vimentin filaments are frequently found in close proximity to FAK-labeled FAs, not only at the leading edge during FA maturation but also at the retracting edge during FA disassembly (Fig. 4A, B; Video 4A_4B). Notably, vimentin contacts were also observed at putative nascent adhesions, identified as small, peripheral FAK-positive structures lacking the elongated morphology of mature FAs (Fig. S4 B), underscoring the broad spatial engagement of vimentin with FAs across the entire cell footprint.

**Figure 4.**
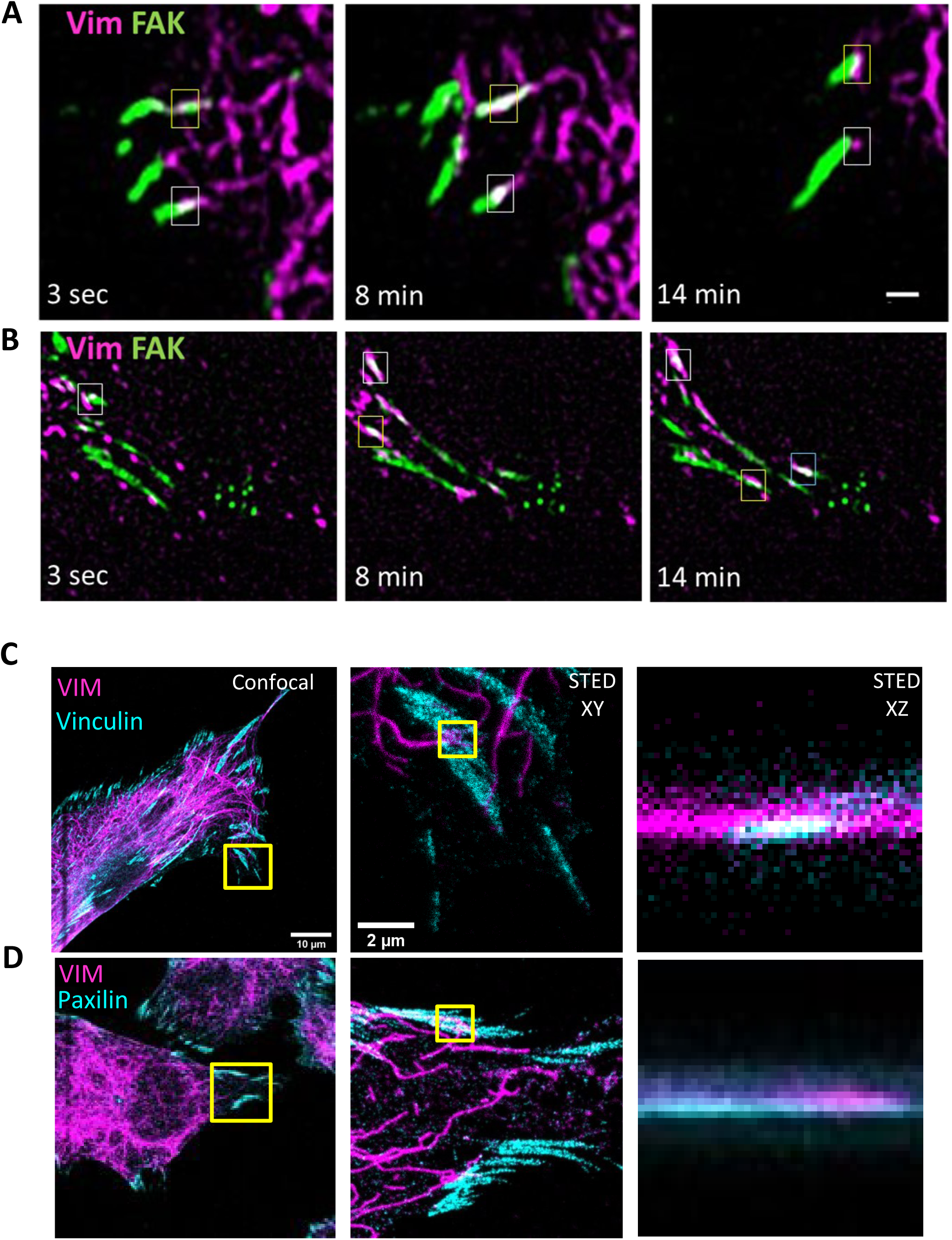
Vimentin interacts with maturing and retracting focal adhesions. (A–B) Time-lapse images of Vim^−/−^ MEFs co-expressing mCherry-vimentin (pseudo-colored in magenta) and GFP-FAK (green), showing interactions with maturing and retracting FAs. (C–D) Endogenous vimentin colocalization with vinculin or paxillin in WT MEFs imaged by confocal and STED microscopy. Insets show XY and XZ views of individual adhesions. Scale bar, 1 µm.

**Figure 5.**
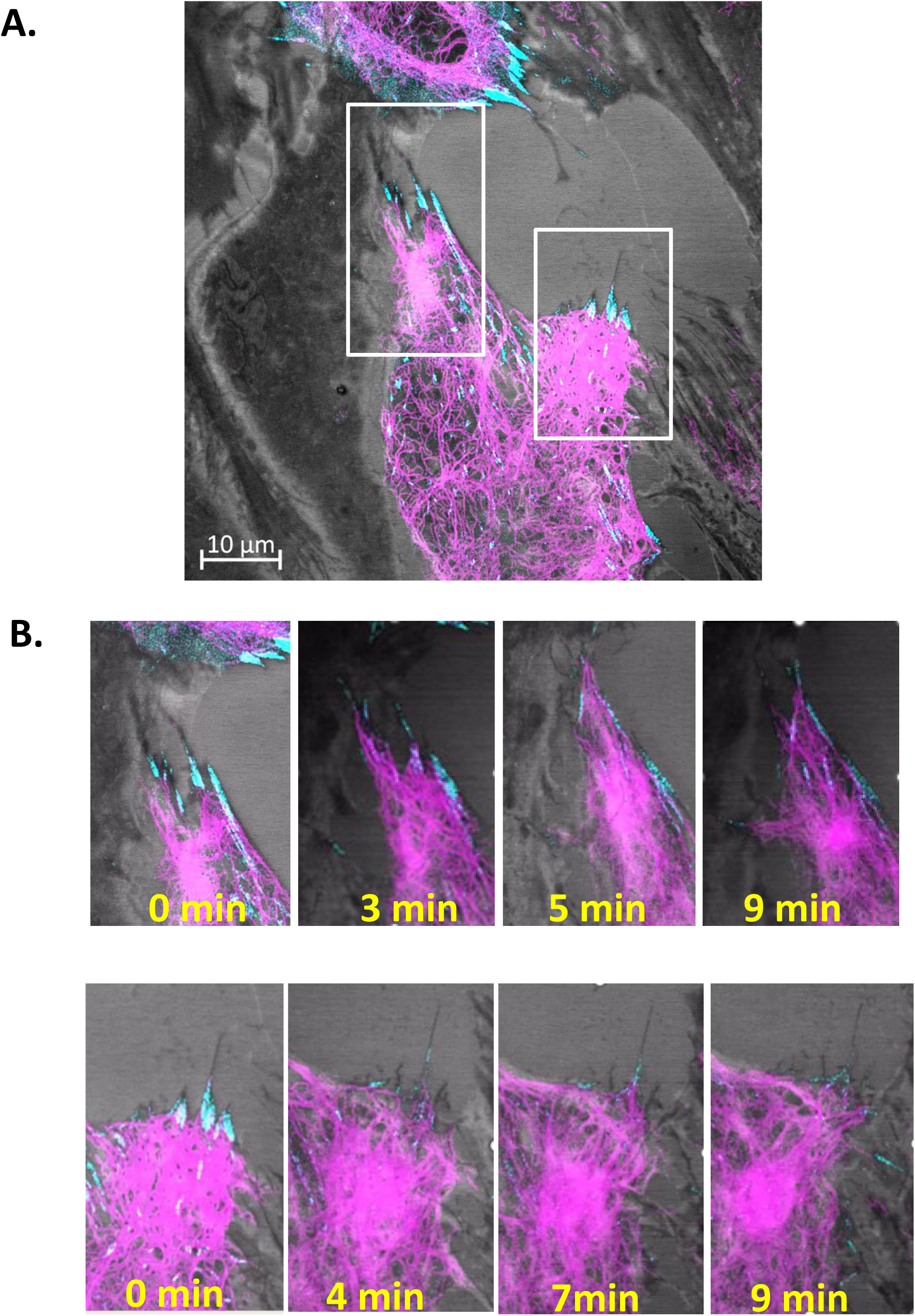
Vimentin pulls on focal adhesions via filament coiling to enable FA displacement. (A) Representative frame from live-cell imaging of MEF vimentin-KO cells expressing mCherry–vimentin (pseudo-colored in magenta) and FAK-GFP-labeled focal adhesions (pseudocolored in cyan). White boxes indicate regions shown at higher magnification in (B). (B) Time-lapse sequences from the boxed regions showing vimentin filament coiling and retraction at adhesion sites, accompanied by focal-adhesion displacement over time. Images were acquired every 2 min for 2–3 h using Airyscan super-resolution microscopy (Zeiss LSM 880, 63× oil objective). Scale bar: 10 µm.

To further elucidate the nature and spatial configuration of vimentin-FA interactions, we employed 2D and 3D STED super-resolution microscopy to visualize endogenous vimentin filaments in relation to key FA proteins. Super-resolution imaging and quantitative image analysis revealed consistent three-dimensional correspondence, such that vimentin-FA overlap observed in the XY plane was maintained along the Z-axis, indicating spatially aligned interactions with multiple FA components, including vinculin (Fig. 4C), paxillin (Fig. 4D), and autophosphorylated FAK (pFAK, Fig. S4A – B).

Phosphorylated FAK was examined specifically because it marks catalytically active, force-engaged adhesions undergoing signaling-dependent maturation and turnover. Notably, vimentin filaments were observed to project toward and insert into adhesion sites, often terminating at or within the FA plaque region. This was evident both in the XY plane 2D STED (Fig. S4A,B) and in vertical 3D STED XZ sections (Fig. S4A,B), where the Z-stack reconstructions revealed a striking vertical alignment of vimentin filaments and phospho-FAK puncta. This spatial proximity strongly supports a model in which vimentin IFs form a close physical interface with the FA complex.

Furthermore, the 3D STED images (Fig. S4 A, B) show that vimentin not only approaches the plane of the substrate but often aligns with phospho-FAK-rich structures in a defined stratified architecture. These interactions appear selective and spatially organized, suggesting that vimentin may target specific subsets of FAs, particularly those associated with mechanical signaling and cytoskeletal remodeling. The localization of vimentin at sites enriched in phospho-FAK, vinculin and paxillin also raises the possibility that vimentin contributes to regulating FA maturation and force transduction at the molecular level.

These dynamic and super-resolution data indicate that vimentin is not merely peripherally associated with FAs but forms intimate, three-dimensional contact zones with their core components. This supports a model in which vimentin directly engages with the adhesion machinery to spatially organize and stabilize FAs, thereby contributing to the maintenance of adhesion integrity and polarized migration. Related to this model, the observations spatial distribution of vimentin-FA contacts, including at retracting edges, is particularly significant given the critical role of trailing edge adhesions in maintaining global traction force balance. As effective directional migration depends on coordinated forces at both protruding and retracting regions, these findings suggest that vimentin contributes to the tuning of net force balance required for persistent cell polarity and migration.

### Optical flow analysis reveals coordinated vimentin remodeling associated with FA displacement

Beyond passive colocalization or structural support, vimentin exhibit dynamic behaviors that appear to actively modulate FA positioning and turnover. To explore these dynamics, we conducted live-cell imaging of migrating cells expressing fluorescently tagged vimentin and FA components.

Strikingly, thick vimentin bundles near the nuclear region often underwent coiling or contraction-like motions that propagated along thinner filaments extending toward the leading edge, mechanically pulling distal FAs toward the cell center ( 5B, Video 6B_6C). In the top time-lapse sequence (0–9 min), this centripetal movement was evident as peripheral FAs positioned at the tips of elongated vimentin filaments were progressively drawn inward during filament retraction(5A, Video 6A). In the bottom sequence, a broader vimentin meshwork displayed lateral sliding and reorganization, accompanied by synchronous displacement and partial disassembly of nearby FAs. Higher-resolution time-lapse analyses of vimentin-FA co-dynamics confirmed that these filament rearrangements coincide with changes in FA morphology and position over time. Collectively, these coordinated behaviors demonstrate that vimentin is not a passive structural scaffold but an active mechanical element capable of modulating adhesion positioning through dynamic filament contractions and lateral translocations, thereby influencing the cytoskeletal force balance during migration.

Importantly, these movements were not stochastic but occurred in coordinated spatial and temporal patterns. In several cases, the coiling of vimentin filaments preceded and appeared to drive the movement of FAs, supporting a model where internal vimentin dynamics exert pulling forces on peripheral adhesions. It is attractive to consider that this mechanical action could function to recycle or reposition FAs during retraction phases of migration or cytoskeletal remodeling.

To rigorously quantify the complex cytoskeletal behaviors described above, we applied optical flow analysis to super-resolution time-lapse sequences. This approach treats the cytoskeleton as a flowing continuum, allowing us to mathematically decompose its movement dynamics into three physical behaviors: speed, contraction, and coiling. First, we calculated the instantaneous velocity magnitude (|v| = √(u² + v²)) to determine the nature of the motion (Fig. S5A). Given the high spatial resolution (0.0425 µm/pixel), this analysis revealed that vimentin networks exhibit slow, persistent retrograde velocities, typically fluctuating between 0.001 and 0.004 µm/s (0.06–0.24 µm/min). This slow crawl is distinct from rapid motor-driven cargo transport (∼1.0 µm/s) (Ross et al., 2008), confirming that the network is not trafficking vesicles but rather undergoing elastic cortical flow and remodeling (Robert et al., 2019; Hookway et al., 2015).

To determine how the network reshapes itself during this flow, we measured the Divergence (∇·v = ∂u/∂x + ∂v/∂y), which quantifies whether the meshwork is actively shrinking (convergence) or expanding (divergence) (Fig. S5B). Our analysis showed that mean flow divergence remained remarkably low (± 5 × 10^−6^ s^−1^), fluctuating weakly between net convergence and expansion. This suggests that the flow does not result in significant isotropic contraction. Instead, to capture the complex reorganization of the filaments, we calculated the Curl (∇×v = ∂v/∂x – ∂u/∂y), which measures the rotational component of the flow field (Fig. S5B). Strikingly, rotational components (|curl|) were consistently an order of magnitude higher than divergence (∼3 × 10^−5^ s^−1^). This indicates that the vimentin flow pattern is dominated by local rotational movements and twisting rather than simple compression.

We further quantified the coordination of this motion using circular variance, where a value of 0 implies perfect alignment and 1 implies chaotic motion (Fig. S5C). The data revealed large dynamic fluctuations, cycling between phases of highly aligned motion (variance ∼0.09) and disordered reorganization (variance > 0.8), indicating that remodeling occurs as intermittent “bursts” of coordination. Finally, to determine how these vimentin flow dynamics affect FA movement, we performed dual-channel flow analysis on FAK (Fig. S5A). We observed a strong positive correlation (Pearson’s r = 0.76) between vimentin remodeling speed and FAK displacement speed. Major bursts in vimentin flow consistently coincided with peak FA sliding events, suggesting that the dynamics of the intermediate filament network are tightly coupled to the regulation of FA mobility.

Taken together, the live-cell data presented in the included videos raise the possibility that coiling, shortening, and lateral translocation of vimentin filaments act in concert with other cellular force-generating and force-bearing systems and signaling-dependent regulatory processes to transmit and redistribute mechanical forces locally. This cooperative coupling likely contributes to the repositioning of FAs. Based on all results above, this cooperative mechanical coupling is likely to facilitate efficient FA turnover, redistribution, or recycling, enabling the spatial plasticity required for persistent and directional migration

### Vimentin dynamically and periodically associates with selective focal adhesions during migration

The analysis of individual FAs described indicated that interactions between vimentin and FAs may take place in a coordinated fashion. Therefore, we wanted examine whether the physical interactions occur in a temporally coordinated manner at the cellular level and also how vimentin influences FA dynamics at a FA population level. To this end, we performed high-resolution time-resolved analyses of vimentin-FA dynamics in migrating cells by live-cell imaging of cells co-expressing eGFP-FAK (green) and mCherry-vimentin (magenta) (Video 7). Strikingly, these analyses revealed that the movements described above were not stochastic but organized in coherent spatial patterns and synchronized over time at the cellular scale (Video 7). Notably, vimentin interactions with FAs were neither continuous nor uniformly distributed across the cell. Instead, the spatiotemporal dynamics of vimentin-FA coupling displayed a periodic pattern of engagement and disengagement. Frame-by-frame colocalization analysis revealed rhythmic fluctuations in spatial overlap between vimentin and FAs, characterized by alternating phases of high and low correlation occurring on the order of several minutes, a timescale consistent with known cycles of FA maturation and turnover (Fig. S6A).

To define the subsets of vimentin IF structures spatially associated with FAs, we performed centroid-based distance tracking between individual vimentin assemblies and focal adhesions over time, allowing quantification of their proximity distribution. This analysis revealed that a distinct fraction of vimentin assemblies maintained persistent spatial proximity to FAs. (Fig. S6B).

Taken together, the spatio-temporal analysis of vimentin-FA interactions revealed that vimentin does not uniformly associate with FAs but instead engages with a selective, spatially and temporally restricted vimentin IF population during defined temporal windows, consistent with most vimentin IFs being distributed throughout the cytoplasm rather than at FAs. This selective spatiotemporal association establishes a framework for dissecting the nature of specific vimentin-FA interactions.

To dissect the heterogeneity of vimentin-FA interactions in greater detail, we performed a high-dimensional analysis of 481 individual FA trajectories. We extracted a seven-dimensional phenotypic signature for each track, comprising instantaneous velocity, directionality, Mean Squared Displacement (MSD) exponent, FA area, Pearson correlation coefficient (PCC), and fluorescence intensities for both FAK and vimentin. Unsupervised Gaussian Mixture Modeling (GMM) segregated the adhesion tracks into three statistically distinct subpopulations based on these multiparametric signatures.

To visualize the temporal evolution of these phenotypes, we first analyzed the protein intensity dynamics over the adhesion lifetimes. This revealed striking differences in composition stability. Cluster 1 (orange) maintained persistently high vimentin levels throughout its lifetime (Fig. 6A), whereas Cluster 3 (green) exhibited low vimentin but high, fluctuating FAK levels (Fig. 6B). This dynamic separation was confirmed by quantitative analysis of the mean intensities, which showed that Cluster 3 is significantly enriched in FAK (Fig. 6C), while Cluster 1 is defined by peak vimentin recruitment (Fig. 6D). Dimensionality reduction via Principal Component Analysis (PCA) confirmed that these clusters represent discrete phenotypic states rather than a continuous distribution (Fig. 6E), driven by distinct structural and motility parameters (Fig. S7E, S7F). To understand the biochemical basis of these states, we examined the relationship between the two major protein markers. A scatter plot of mean vimentin versus mean FAK intensity revealed a non-linear distribution, indicating that FAK accumulation and vimentin recruitment are decoupled events (Fig. 6F).

**Figure 6.**
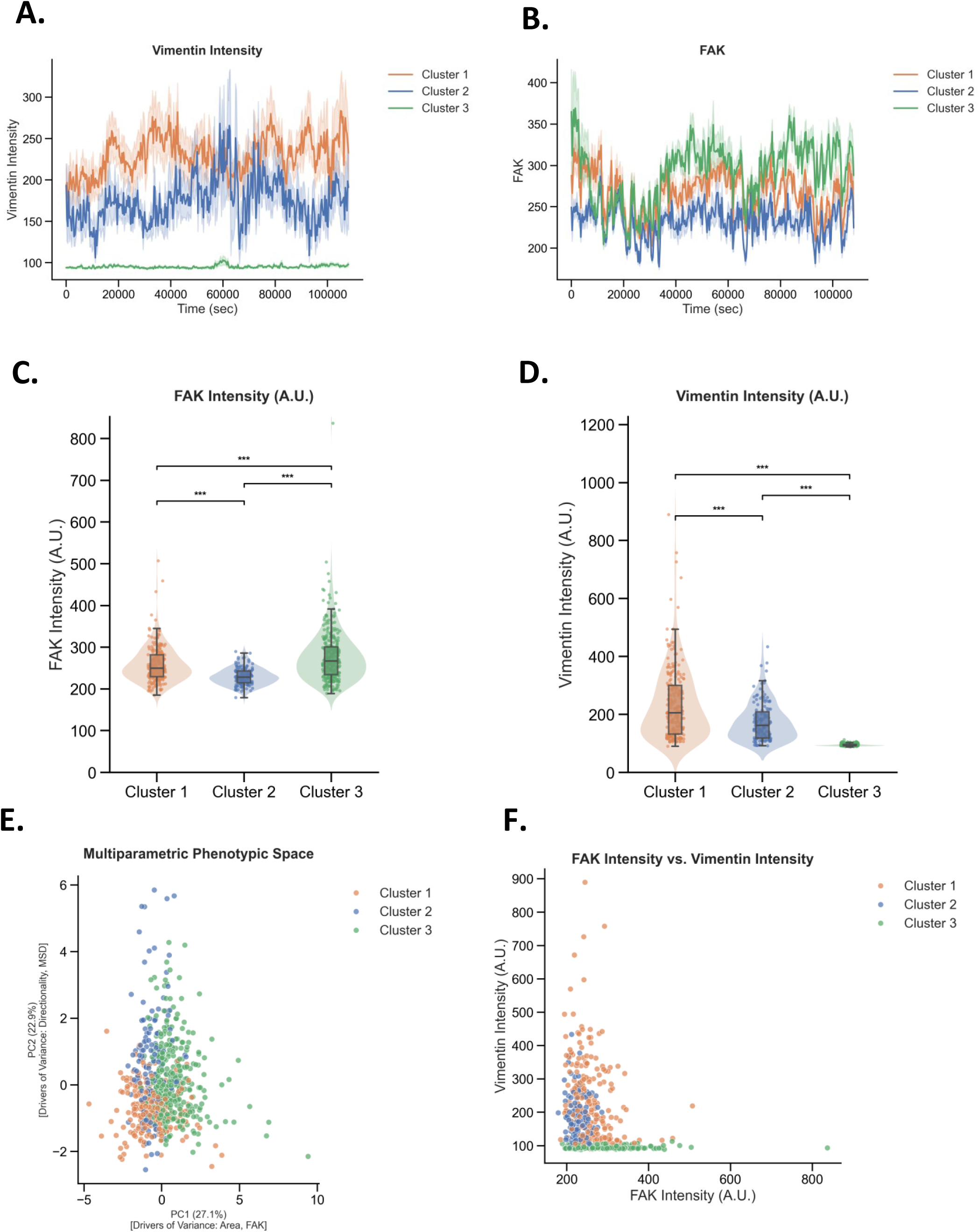
Vimentin recruitment defines focal adhesion maturation states. (A–B) Time-resolved analysis of (A) Vimentin intensity and (B) FAK intensity over the lifetime of adhesions. Solid lines represent the population mean; shaded areas represent SEM. Cluster 1 (orange) maintains high stable Vimentin, while Cluster 3 (green) shows fluctuating high FAK. (C–D) Raincloud plots showing the distribution of (C) Mean FAK intensity and (D) Mean Vimentin intensity across the three clusters. Cluster 3 is FAK-dominant; Cluster 1 is Vimentin-dominant. (E) Principal Component Analysis (PCA) of the 481 focal adhesion trajectories. The distinct separation of clusters (colors) confirms they represent discrete phenotypic states. (F) Scatter plot of mean Vimentin versus mean FAK intensity. The lack of a linear correlation demonstrates that FAK accumulation and Vimentin recruitment are independent, decoupled processes.

Cluster 1 represents a mechanically anchored state. Beyond its high vimentin content (Fig. 6A, 6D), this population displayed the lowest MSD values (Fig. S7A) and the lowest directionality (Fig. S6B), indicating a high degree of spatial confinement and minimal net displacement. The convergence of high vimentin load with near-zero directionality suggests that heavy vimentin recruitment acts as a mechanical scaffold that effectively traps the adhesion. Temporal analysis confirmed that this high-vimentin, low-mobility state remains stable over the adhesion lifetime, identifying Cluster 1 as mature, load-bearing traction points(Fig. 6A).

Cluster 2 represents a transient, dynamic state. Characterized by the lowest mean FAK intensity (Fig. 6C) and intermediate vimentin levels (Fig. 6D), these tracks exhibited broad MSD distributions (Fig. S7A) consistent with exploratory or oscillatory motion. The low abundance of signaling proteins combined with fluctuating motion suggests these adhesions are weakly engaged with the extracellular matrix and likely participate in adhesion turnover or leading-edge sampling.

Cluster 3 represents a signaling-competent but mechanically unconfined state. Despite significant enrichment of FAK (Fig. 6C), these adhesions did not exhibit the spatial confinement observed in Cluster 1. Instead, Cluster 3 tracks remained mobile, with MSD values significantly higher than the vimentin-rich Cluster 1 (Fig. S7A). Furthermore, the correlation between vimentin and FA signals (PCC) was consistently lower in this group compared to Cluster 1, both in mean distribution (Fig. S7C) and over time (Fig. S7D). This decoupling reveals that high FAK content alone does not confer mechanical anchoring; rather, these adhesions distinguish biochemical enrichment from mechanical maturation.

Collectively, these data demonstrate that FA maturation segregates into discrete functional states. While FAK accumulation marks the biochemical maturation of the adhesion (Cluster 3), it is the specific recruitment of vimentin that correlates with the transition to a spatially confined, mechanically anchored state (Cluster 1). These results support a model where vimentin acts as a mechanical brake, stabilizing focal adhesions against motility only after they have been established.

Strikingly, these results reveal that vimentin engages with FAs in a temporally patterned and subpopulation-specific manner. Rather than forming static scaffolds, vimentin filaments dynamically interact with a defined subset of stable, anchoring adhesions in a periodic cycle that mirrors FA maturation and disassembly. This selective reinforcement likely plays a critical role in maintaining directional persistence during migration by stabilizing mechanical feedback at specific adhesion sites. The loss of such targeted anchoring in vimentin-deficient cells provides a plausible mechanistic basis for the disorganized FA dynamics and impaired migratory behavior observed earlier (Fig. 1), positioning vimentin as a spatiotemporal integrator of adhesion stability and cytoskeletal coordination.

### Vimentin is a dynamic component of the FA nanoarchitecture

Since our live-cell observations and STED super-resolution data revealed dynamic interactions between vimentin filaments and FAs, we next wanted to examine in detail whether vimentin is structurally integrated within the nanoscale architecture of FAs as described previously (Kanchanawong et al., 2010). STED microscopy, while powerful in lateral resolution, lacks sufficient axial precision to resolve this question. Therefore, we turned to interferometric Photoactivated Localization Microscopy (iPALM), which provides ∼20 nm isotropic resolution in all three spatial dimensions, enabling nanoscale mapping of vimentin’s 3D spatial relationship with FA proteins, using representative x-y and x-z projections (Fig. 7A-C) and subsequent quantitative z-center analysis.

**Figure 7.**
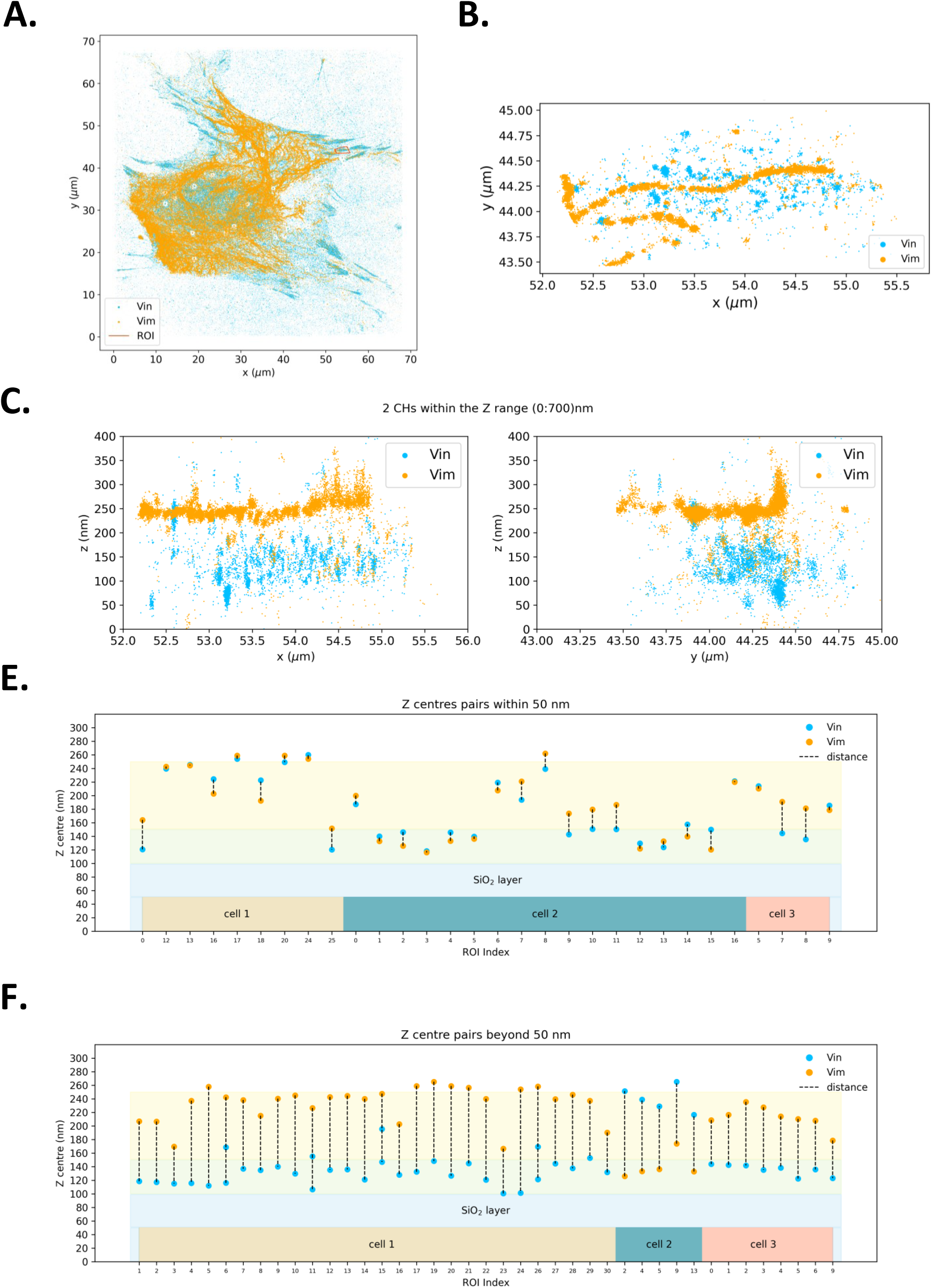
Vimentin is a dynamic component of the focal adhesion nanoarchitecture. (A–F) iPALM analysis of vimentin and vinculin localization across FA layers reveals that vimentin spans multiple FA protein strata and dynamically associates with the force-transduction region.

To quantify axial localization, we calculated z-centers, the peak vertical positions of vimentin and vinculin (a core mechanotransductive FA protein) localizations, by fitting iPALM-derived point cloud data with Gaussian or double-Gaussian models (Fig. 7A-F). In several adhesions, vinculin exhibited bimodal z-distributions, allowing subpopulation analysis that revealed vimentin signals either overlapping with or positioned above the upper vinculin band. These reference examples (Fig. S8) illustrate the fitting approach used for all quantitative analyses.

Using iPALM in TIRF mode to restrict imaging to the basal adhesion interface (0-700 nm above the coverslip), we consistently observed vimentin localizing within the vertical axis of FAs, occupying axial positions between 100 and 260 nm (Fig. 7E-F, showing z-center pairs within ≤50 nm or >50 nm separation, respectively). While not every FA contained detectable vimentin signal, the majority did, suggesting that vimentin integration into FAs may be transient or context-dependent at the level of individual adhesions, and consistent with its dynamic behavior observed in time-lapse imaging with GFP-FAK (Fig. 4A-B; Video 4, 5,7).

Single-molecule localization microscopy using iPALM revealed that vimentin molecules are closely associated with vinculin, a core mechanotransductive component of FAs, but display substantial axial heterogeneity (Fig. 7E-F, Video 8). In most adhesions, vimentin overlapped or was positioned slightly above vinculin along the z-axis, whereas in others, dual vimentin populations were evident, one coinciding with the vinculin layer and another situated above it. This variation indicates that vimentin is not restricted to a single structural plane but spans multiple FA strata, including regions corresponding to the force-transduction and actin-regulatory layers described by Kanchanawong et al. (2010), where force-dependent conformational changes in vinculin and talin are known to occur (Case et al., 2015; Sun et al., 2016).

To quantitatively assess these distributions, we determined the z-center positions of vinculin and vimentin by Gaussian or double-Gaussian fitting of iPALM localization data from 58 individual adhesions (Fig. S9). In many cases, vinculin exhibited bimodal vertical distributions, whereas vimentin was more broadly distributed, often overlapping both vinculin peaks. Across all adhesions, the mean z-center for vinculin was 158.9 ± 45.4 nm and for vimentin 205.6 ± 43.5 nm, consistent with vimentin occupying higher layers on average. Importantly, the substantial spread in z-center values for both proteins indicates that neither vinculin nor vimentin occupies a fixed axial position across adhesions, but instead displays pronounced nano-scale variability at the level of individual FAs. However, the partial overlap of both proteins and the occurrence of dual z-populations for each (single– and two-peak fits; Fig. S8A-F) indicate that vimentin associates with multiple nano-layers within the adhesion body, consistent with its dynamic and context-dependent engagement observed in the live-cell analyses described above.

The majority of vimentin-vinculin pairs resided within ∼50 nm of each other along the z-axis, consistent with a close structural association, while a subset showed larger offsets (>100 nm), reflecting variable or transient alignment. This axial heterogeneity is fully consistent with the dynamic behaviors observed in live-cell imaging and FA clustering analyses described above, in which vimentin engagement with adhesions was heterogeneous, temporally regulated, and dependent on adhesion state. The observed heterogeneity likely reflects the dynamic nature of vimentin integration into FAs, consistent with time-lapse imaging showing vimentin filament coiling and retraction at adhesion sites (Fig. 5 A-B; Video 6A). Together, these data demonstrate that vimentin is not merely juxtaposed to FAs but is incorporated into their nanoarchitecture, spanning multiple functional layers where it may modulate mechanotransduction and adhesion stability.

Together, these iPALM data demonstrate that vimentin is not merely adjacent to FAs but is integrated within their nanoarchitecture in a context-dependent manner. Its axial positioning alongside vinculin within defined FA layers supports a model where vimentin participates in mechanotransduction and contributes to adhesion stability and force transmission from within the adhesion complex itself. Rather than adopting a single, invariant axial position, vimentin appears to dynamically engage distinct FA nano-layers in a manner that parallels its mechanically and temporally regulated behavior observed at the cellular scale.

### Integrative interactome analysis supports vimentin association with focal adhesion components

Previous work from our group established vimentin as a multifunctional signaling scaffold capable of interacting with diverse regulatory proteins (Pallari & Eriksson, 2006). As part of a broader effort to systematically identify vimentin-associated protein networks across cellular contexts, we have used affinity purification-mass spectrometry (AP-MS) to characterize vimentin interactomes in multiple experimental settings, most recently including nutrient-sensing pathways and mTORC1 signaling (Mohanasundaram et al., 2022). In the present study, we analyzed a vimentin AP-MS interactomics dataset to determine whether vimentin-associated proteins include components relevant to FA organization and cytoskeletal regulation. The dataset used for this analysis is described in the Methods and the accompanying tables that have been uploaded for open access (see Methods section for details).

To test whether the FA-associated properties and behaviors of vimentin observed in our imaging experiments are reflected at the molecular interaction level, we analyzed high-confidence vimentin interactors identified by AP-MS (MiST ≥ 0.93), derived from datasets generated under nutrient-rich and starved conditions.This analysis allowed us to assess whether proteins known to occupy distinct structural layers of focal adhesions are represented within the vimentin-associated interactome.The analysis revealed multiple proteins with established roles in FA assembly, actin regulation, and cytoskeletal signaling, indicating that vimentin associates broadly with proteins positioned at the adhesion-cytoskeleton interface (Fig. S10; Table 1).

**Table 1:** Vimentin interactors relevant to focal adhesion organization and actin remodeling.

| Functional module | High-confidence interactors (Gene Name) | Principal function / rationale for inclusion |
| --- | --- | --- |
| Actin remodeling & FA assembly | ENAH, FMN2, CAPZA1, ACTN4, DBN1 | Drive actin filament elongation, capping, and bundling at adhesion sites; regulate lamellipodial protrusion and adhesion turnover. |
| Integrin-mediated signaling | TLN1, TNS1, ZYX, FLOT2, EMILIN1, NID2 | Couple integrins to the actin cytoskeleton and ECM; mediate adhesion signaling, integrin clustering, and caveolar trafficking. |
| Force transmission & contractility | VASP, MYLK, ACTL6A, CNN2 | Link vimentin to actomyosin tension and FA mechanosensing; control contractile feedback and adhesion maturation. |
| Cytoskeletal regulation / mechanotransduction | KRT18, LUZP1, SCYL2, SMARCE1 | Structural or scaffolding roles supporting cytoskeletal integrity and signaling; potential indirect links to vimentin filament organization. |
| Transcriptional / signaling adaptors (contextual) | EP300, CREBBP, ARID1A | High-score nuclear interactors possibly reflecting mechanosensitive signaling through vimentin scaffolds; listed for completeness but not used in FA modeling. |

Functional annotation of these interactors revealed four modules relevant to FA organization:

Proteins involved in **actin remodeling and FA assembly** were prominent, including ENAH, FMN2, CAPZA1, ACTN4, and DBN1. These proteins regulate actin filament elongation, capping, and bundling at adhesion sites and support lamellipodial dynamics and adhesion turnover, processes known to be disrupted in vimentin-deficient cells.

Several **integrin-associated adhesion regulators** were identified, including TLN1, TNS1, ZYX, FLOT2, EMILIN1, and NID2. These molecules contribute to integrin-cytoskeleton coupling, integrin clustering, and mechanosensitive signaling within FAn complexes.

The interactome included **regulators of actomyosin force transmission and contractility**, such as VASP, MYLK, and CNN2, together with signaling regulators including ACTL6A. These proteins participate in actin filament elongation, myosin activation, and cytoskeletal tension generation, processes essential for FA maturation and force transmission.

A set of **cytoskeletal regulators and contextual mechanotransducers**, including KRT18, LUZP1, SCYL2, and SMARCE1, may reflect broader structural or regulatory connections between intermediate filament networks and adhesion-associated signaling pathways.

These molecular associations outline a hierarchical organization consistent with the nanoscale architecture of FAs described by super-resolution imaging studies. In this framework, FAs are organized into vertically stratified layers linking integrin-based matrix adhesion to actin assembly and cytoskeletal force transmission.At the cell-matrix interface, extracellular matrix engagement is transmitted through integrin complexes that couple to FA scaffolds such as talin (TLN1) and zyxin (ZYX). Downstream of these complexes, actin assembly regulators including FMN2, ACTN4, and VASP promote filament nucleation, elongation, and bundling at adhesion sites. These actin structures are linked to contractile machinery through regulators such as MYLK and CNN2 that control actomyosin-generated tension and FA maturation. Within this framework, the vimentin IF network occupies a strategic position downstream of actomyosin-generated tension, where it can integrate mechanical forces generated at adhesions and coordinate their propagation across the cell.

Importantly, these associations should not be interpreted as evidence of stable, constitutive binary binding interactions but rather reflect the dynamic, multilayered organization of FA complexes and their supporting cytoskeletal systems. FAs are highly dynamic macromolecular assemblies composed of hundreds of proteins that continuously assemble, remodel, and disassemble in response to mechanical and signaling cues. In this context, proteins identified in vimentin-associated interactomes likely represent components of transient, context-dependent molecular environments rather than fixed binding partners. Consistent with this view, our imaging analyses show that vimentin filaments associate dynamically with FAs, selectively engaging subsets of adhesions and exhibiting temporally regulated interactions during adhesion maturation, displacement, and disassembly. These observations indicate that vimentin participates in a fluctuating interaction landscape in which cytoskeletal and adhesion components are recruited, reorganized, and released over time, such that the interactors identified here likely represent proteins coexisting within shared mechanical and signaling microenvironments at the adhesion-cytoskeleton interface.

Within this framework, vimentin-associated proteins span several functional levels of the adhesion system: (i) actin regulators influencing adhesion formation and turnover, (ii) mechanosensitive FA components participating in force transmission, and (iii) signaling regulators modulating cytoskeletal tension and downstream feedback pathways. Together, these layers form a hierarchical axis linking extracellular matrix engagement through integrins and FA scaffolds to actin assembly, actomyosin contractility, and the vimentin intermediate filament network.

Because these interactions occur within a highly dynamic and spatially heterogeneous system, definitive validation of individual molecular interactions, including identification of binding interfaces, interaction motifs, and stoichiometric relationships, would require extensive biochemical and structural analyses beyond the scope of this study. Instead, the strength of the present proteomics approach lies in integrating proteomic interaction landscapes with live-cell and super-resolution imaging to define the systems-level organization in which vimentin operates.

Taken together, the proteomic and imaging data support a model in which vimentin participates in a modular, temporally regulated cytoskeletal network coordinating adhesion assembly, force generation, and directional migration. In this model, vimentin functions not as a static adhesion component but as a responsive mechanical scaffold integrated within the broader adhesion machinery.

### Vimentin guides FAs to maintain the directional persistence

Taken together, our findings position vimentin as a dynamic and mechanosensitive integrator within the adhesion-cytoskeleton system, essential for coordinated cell migration. In fibroblasts, vimentin filaments align with the cell’s front-rear polarity axis and dynamically associate with FAs, where they undergo contraction-like motions that physically reposition adhesions and promote directional migration (Fig. 8, top left). In contrast, vimentin-deficient cells lose this organized filamentous architecture, resulting in dispersed, misoriented adhesions and unstable migration trajectories (Fig. 8A, top right). Super-resolution imaging and iPALM nanoscopy further revealed that vimentin is not merely peripherally associated with FAs but is embedded within their vertical nanoscale structure (100–260 nm), overlapping the actin-regulatory and force-transduction layers (Fig. 8B). In this position, vimentin filaments interface with core components including vinculin, talin, and FAK supporting mechanical feedback and maturation of adhesions.

**Figure 8.**
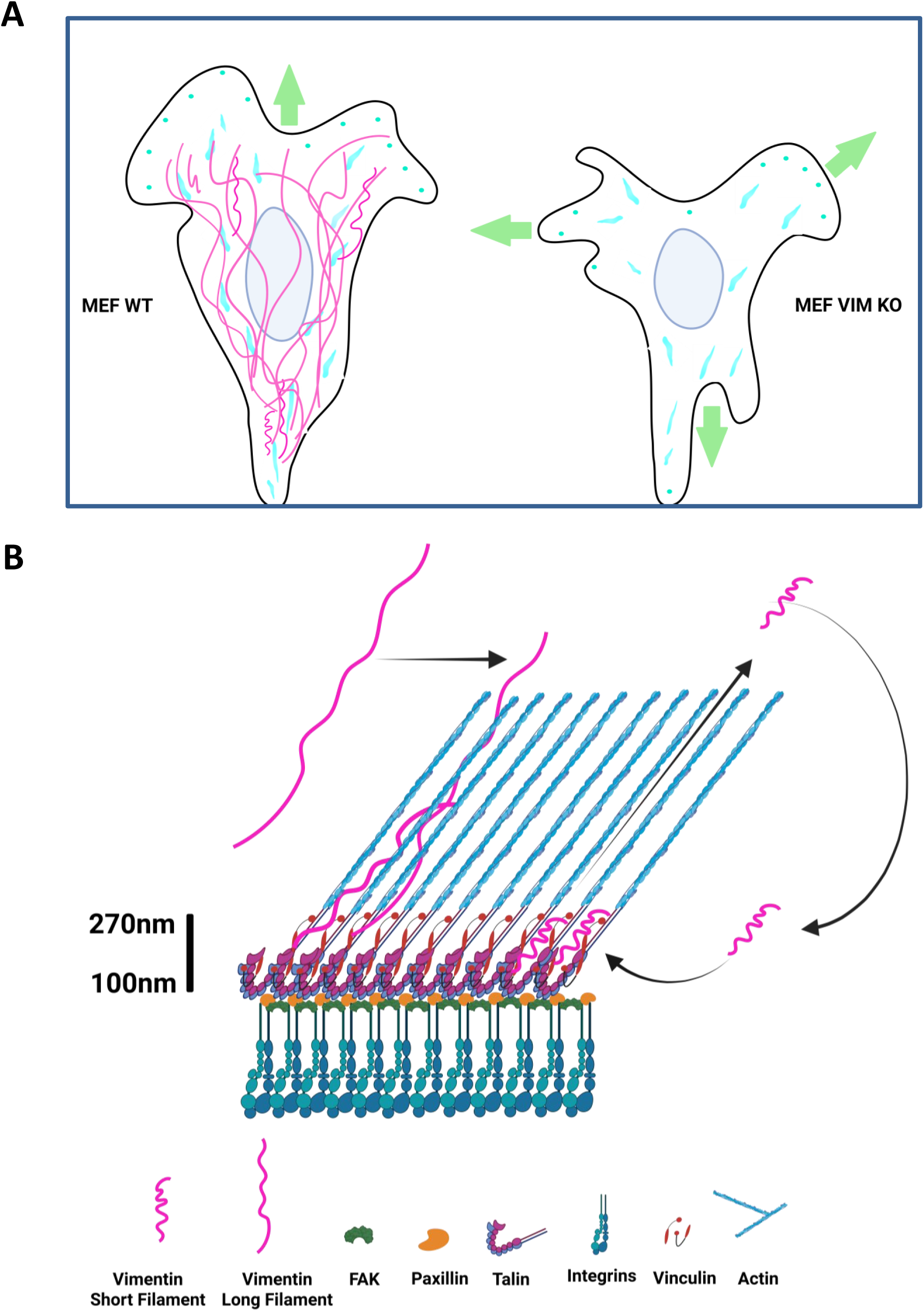
Model: Vimentin steers focal adhesions to maintain directional persistence. Proposed model illustrating how vimentin regulates FA orientation, size, and spatial organization to buffer actomyosin-generated traction forces. Vimentin–FA interactions stabilize adhesion positioning and alignment, enabling sustained directional migration. Loss of vimentin disrupts FA coordination, resulting in irregular force transmission and loss of directional persistence.

Integrative proteomics confirmed this architectural integration, identifying high-confidence vimentin interactors across multiple FA modules: actin-remodeling proteins (e.g., ENAH, FMN2), integrin-associated linkers (e.g., TLN1, TNS1, ZYX), and force-transmission effectors (e.g., VASP, MYLK). The iPALM observations together with the findings from interactomics suggest that vimentin participates in a multilayered FA network: at the base, by linking to actin regulators driving adhesion growth and turnover; within the force-transduction layer, by interacting with mechanosensitive scaffolds; and at higher levels, by engaging components modulating contractility and cytoskeletal signaling. This multi-tiered integration enables vimentin to regulate FA stability, alignment, and mobility in response to mechanical cues, acting as both a scaffold and an active force transmitter. Altogether, vimentin supports a feedback loop that aligns FA dynamics with cellular polarity, thereby sustaining persistent migration in mesenchymal cells

Our results show that vimentin acts to coordinate the FA orientation to yield directional migration of fibroblasts. The assumed modus operandi suggests coordination of an FA-based all-wheel drive of migrating fibroblasts to maintain directional persistence and as well as high levels of directionality (Fig. 8B). Previous studies show that vimentin IFs interact with actomyosin complexes of the cell migration machinery (Jiu et al., 2015, 2017) and here we show that vimentin resides and operate dynamically in close proximity to critical FA protein vinculin suggesting vimentin resides around force transduction layer of FA nanostructure. In summary, the bidirectional interaction of vimentin with, as previously shown, the actin arcs and stress fibers (Jiu et al., 2015, 2017) and, in this study, with the FA complexes, yields the proper contractile and traction forces as well as coordination of the FAs that is required for the net directional movement of a migrating fibroblast.

## Discussion

The mechanisms through which vimentin controls cell migration and especially cell directionality is still obscure. Here, we show that vimentin is required for directional persistence and that both long filaments and short filaments interact with FAs in a way that determines their dynamics, spatial organization, stability, and orientation in migrating cells. Vimentin squiggles move in a retrograde flow towards the rear end of the cell to interact with the FAs and in the absence of vimentin, the average size of FAs is dramatically reduced. Together, our results reveal a role for vimentin as a regulator of the dynamics and localization of FAs and uncover how the interplay between vimentin and FAs control directional migration of fibroblasts.

Our findings provide a mechanistic framework linking intermediate filaments to adhesion-based force coordination during migration. Previous studies of collective migration have shown that disruption of the intermediate filament network alters force transmission and directional coherence (De Pascalis et al., 2018). In 3D environments, vimentin has also been implicated in regulating matrix adhesion and traction forces (Rodriguez et al., 2026). However, how vimentin interfaces with FAs to organize directional migration at the single-cell level has remained unclear. Our data indicate that vimentin does not simply modulate force magnitude but instead organizes the spatial and temporal coordination of FAs. By stabilizing and aligning specific adhesion subpopulations, vimentin converts local adhesion dynamics into persistent directional migration.

The longer lifespan of vimentin IFs has been proposed to provide a basis for a cellular “memory” that maintains polarity in migrating cells (Gan et al., 2016). Vimentin controls cell polarity also by helping to orient and redistribute the traction forces generated by the contraction of actomyosin bundles (Costigliola et al., 2017), a function essential for mechanical resilience and signaling in physiologically demanding contexts (van Engeland et al., 2019; Shaebani et al., 2022; Coelho-Rato et al., 2024). FAs have a crucial role in this process since the traction stress is exerted on the extracellular matrix (ECM) via actin stress fibers attached to FAs. FAs consist of more than 150 components (Zaidel-Bar et al., 2007). They are indispensable for signaling, cytoskeletal reorganization, and force transduction from and to ECM (Kanchanawong et al., 2010). Some preceding studies have indicated vimentin to be recruited to a subset of FAs via plectin1f linker protein (Burgstaller et al., 2010). The vimentin-associated FAs have been shown to exhibit higher adhesion strength and this is mediated by the presence of β3 integrin which is crucial for recruitment of vimentin to the FAs (Bhattacharya et al., 2009). The presence of actin-regulatory and adhesion-associated proteins in the vimentin interactome (Fig. S10) further supports the idea that vimentin resides within a biochemical environment coordinating cytoskeletal remodeling and FA stability.

Vimentin is mechanically and biochemically coupled to FAs and the actin cytoskeleton through linker proteins such as plectin and through interactions with actin-regulatory components (Gregor et al., 2014; Osmanagic-Myers et al., 2015; Jiu et al., 2015). These connections provide a structural basis for integrating intermediate filaments with adhesion complexes and contractile actin networks. While prior studies have implicated these interactions in migration and force regulation, how they contribute to coordinated adhesion dynamics across the cell has remained unclear. Our findings suggest that such coupling enables vimentin to selectively stabilize and align specific adhesion subsets, thereby organizing adhesion field behavior during persistent migration

Related to the loss of FA angular periodicity that we observed, the angular bias of FA formation has been linked to actin network organization and retrograde actin flow, which impose preferred orientations on nascent adhesions during migration (Hotulainen & Lappalainen, 2006; Burnette et al., 2011; Etienne-Manneville, 2013). Given the bidirectional coupling between vimentin and actin systems described above (Jiu et al., 2015; Jiu et al., 2017), the loss of angular periodicity in FA birth observed in Vim^−/−^ cells could be directly related to impaired transmission of actin-derived spatial cues to nascent adhesions.

Our data show that both vimentin long filaments (Fig. 4 A) and short filaments (Fig. 4 B) associate with the maturing and disassembling FAs in anterograde and retrograde directions. These FA-vimentin interactions were not limited to the leading edge of the migrating cell but were also observed in the retracting edges of the cell. Even more intriguing was the way by which vimentin enables the movement of FAs by pulling on them by coiling up into dense structures from loosely located long filaments (Fig. 5, Video 6B_6C). To the best of our knowledge, this kind of interaction and the control of FAs has not been reported elsewhere by an intermediate filament protein. Maintenance of an active control of FA dynamics and orientation across the cell periphery is crucial for balanced distribution of contractile forces created by stress fibers connected to the FAs. Vimentin precursors have been reported to be immobilized by plectin 1f protein in a subset of FAs (Burgstaller et al, 2012) and that this immobilization helps the assembly of long filaments. It is plausible that there is an intermediate process that links the short vimentin filaments to interact with the FAs on cell periphery, to be further transported to the actin transverse arcs. This cycling of vimentin filaments might involve: the immobilization to the FAs, further transport to actin transverse arcs, followed by the retrograde flow to the nuclear periphery.

It important to examine the exact role of vimentin IF networks and their interplay with FA complexes, in cell directionality maintenance in a three-dimensional environment, as well as to uncover the detailed molecular signaling and other mechanisms underlying vimentin-mediated FA formation, disassembly, and the arrangement of FA proteins in different vertical compartments such as integrin signaling layer, force transduction layer and actin regulatory layer. Our effort to resolve this end using iPALM microscopy sheds new light on the active localization of vimentin in the FA nanostructure around the force transduction layer as our data shows (Fig. 7). Existence of vimentin around vinculin also suggests that there could be a potential interaction between talin which is essentially considered to be backbone protein of FAs and a key mechanosensor (Case et al., 2015; Sun et al., 2016).

A major additional finding from our study is the identification of three distinct FA subpopulations through unbiased clustering analysis (Fig. 6). This revealed that vimentin is not uniformly associated with all FAs, but is a key, stable component of a specific subset of slow-moving, “Anchoring FAs.” These vimentin-associated Anchoring FAs exhibit greater stability and are likely central hubs for coordinating FA organization across the cell. Loss of this subpopulation in vimentin-deficient cells disrupts the global coordination of FA dynamics, leading to reduced FA size, increased disassembly rates, and ultimately impaired directional persistence. The periodic fluctuations in vimentin–FA colocalization observed in live-cell imaging suggest that vimentin participates in the cyclical remodeling of adhesions rather than static reinforcement. This dynamic coupling could allow vimentin to synchronize adhesion turnover with actomyosin contractility, maintaining a balance between stability and plasticity during directional movement. This result provides a mechanistic explanation for how vimentin stabilizes and orients FAs in a spatially coordinated manner during migration.

Our data (Fig. 2) suggests that FAs of wild type cells have higher stability than the knockout cells lacking vimentin as supported by previous literature (Tsuruta et al, 2003).While the disassembly rate of FAs in knockout cells was significantly higher, vimentin appears to control FA turnover, which is also supported by the data on the FRAP recovery rate and half time recovery measurements. The faster disassembly rate must be creating inefficient duration for holding the FA components in place to maintain the stability that is required for proper directionality, as clearly indicated by the disorientation of FAs when vimentin is absent. The stability and orientation of FAs go hand in hand (Berginski, M. E., & Gomez, S. M, 2013). Hence, a certain degree of FA stability is required for the spatiotemporal localization of the traction forces generated by the stress fibers connected to the FAs (Costigliola et al, 2017). Our study shows that vimentin has a major impact on these aspects crucial for directional persistence by stabilizing FAs.

While there was no difference in FA numbers between WT and Vim KO fibroblasts, the mean FA area was significantly reduced in Vim KO fibroblasts (Fig. 3C). The reduced area links vimentin to FA stability maintenance. It is tempting to assume that the close interactions with FAK could be related to promotion of the Y397 autophosphorylation site of FAK. This is supported by the previous literature where it was shown that vimentin is required for the activation of FAK through VAV2-Rac1 pathway in lung cancer cells (Havel et al., 2014). Vimentin has also been associated with adhesion regulated signaling as it is crucial for the PKCɛ-mediated β1 integrin recycling that maintains cell adhesion (Ivaska et al., 2005). Vimentin can directly interact with β4 integrin containing adhesions leading to recruitment of Rac1 promoting actin polymerization and directional cell migration (Colburn & Jones, 2018). Vimentin also can bind to the cytoplasmic tail of β3 integrin (Bhattacharya et al., 2009) and might directly affect integrin signaling and the connection to actin and MTs. In addition, vimentin can induce the phosphorylation of the MT-associated GEF-H1 at Ser886 leading to RhoA activation and regulating stress fiber assembly (Jiu et al., 2017). Together, these pathways provide a contextual framework for how vimentin may influence adhesion-dependent signaling without implying that all are directly engaged in the present system.

In summary, our study identifies vimentin as a key organizer of FA coordination required for directional persistence in migrating fibroblasts.Rather than altering focal adhesion assembly per se, vimentin selectively stabilizes, orients, and dynamically coordinates specific FA subpopulations across the cell, thereby synchronizing traction forces at the front and rear. Through this mechanism, vimentin converts local adhesion dynamics into coherent, cell-wide directional migration. We propose a model (Figure 8) in which vimentin functions as an active organizer of FA-based force transmission, enabling directionally persistent migration through an integrated and spatially coordinated “all-wheel drive” mechanism.

## Materials and Methods

### Cell lines, antibodies and plasmid constructs

MEF WT and Vim^−/−^ cells were isolated from wild type mice and vimentin knockout mice respectively in our laboratory (Virtakoivu et al., 2015b). They had previously been immortalized with SV40 transfection lentiviral vector. REF WT and Vim^−/−^ were prepared using the CRISPR Cas9 system as described earlier (Vakhrusheva et al., 2019). All the cells were cultured in DMEM media with high glucose (# D5671, Sigma-Aldrich, USA) supplemented with 10% fetal calf serum (Biowest), 2 mM L-glutamine, 100 U ml-1 penicillin, 100 µg ml-1/ streptomycin. Plasmid constructs of emerald-Paxillin and GFP-FAK were obtained as. mCherry empty vector (EV), vimentin wild-type (WT) and vimentin S71A plasmids were kind gifts from Hong-Chen Chen (Department of Life Sciences and Agricultural Biotechnology Center, National Chung Hsing University, Taichung 402, Taiwan) (Pan, Chen, & Chen, 2011). Antibodies used were: vimentin chicken polyclonal (# 919101, Biolegend), phosho-FAK (Y397) rabbit polyclonal (Cat# 44-624G, Thermo Fischer Scientific), phosho-FAK (Y397) rabbit monoclonal (Cat# ab81298, Abcam), Anti-fibronectin antibody (Sigma-Aldrich, Cat# F3648) Anti-collagen I alpha 1 antibody (Novus Biologicals, Cat# NB600-408), mouse monoclonal vinculin (Sigma Aldrich, Cat# V9131), rabbit monoclonal paxillin (Abcam, Cat# ab32084), rabbit monoclonal FAK (Thermo Fischer Scientific, Cat# 701094), Anti-rabbit Alexafluor488, Anti-rabbit Alexafluor555, Anti-rabbit Alexafluor647, alpha-tubulin rabbit polyclonal (CST), Alexafluor Abberior Star Phalloidin.

### Scratch wound assay

For measuring the speed and directional persistence of the migrating cells, WT and vimentin KO fibroblasts were plated in 6 wells plate (Falcon) till confluency. A scratch was made using 10 µl pipette tip and the cells were allowed to migrate for 24 hrs at 37 C and 5% CO2. The scratch area was imaged every 5 mins using a CellIQ live imaging machine from Chipman Technologies, Finland. The time-lapse Videos obtained were used to track the individual cells for obtaining the single cell trajectories, speed, directionality and directional persistence.

### Transfections

Transient transfections for all the cell lines were performed using Xfect transfection reagent from Takara (Cat# 631318) in 6 well plates (Falcon) as per manufacturer’s recommendations.

Cell Derived Matrix generation and Directional Migration Assay Cell derived matrix was generated from WT MEFs in 24 well plate format as described (Kaukonen et al., Nature Methods, 2017).

### Immunostaining and immunofluorescence

Cells were grown on glass coverslips until confluency. Using a 10µl pipette tip confluent cell layer was scratched and allowed the cells to migrate. Cells were fixed using 4% PFA for 10 min at 4 hr time points. Cells were washed with PBS 3X and then permeabilized with 0.2% Triton X-100 in PBS for 15 min at RT, washed in PBS 3X, blocked with 5% Goat serum in PBS for 1 hour in RT. Then the coverslips were incubated on top of 100ul drops of primary antibody in 5% Goat serum in PBS for 1 hour at RT, dip washed 10X in PBS, incubated on 100ul drops of secondary antibody for 1 hour at RT, dip washed the coverslips using forceps 10X in PBS, rinsed in MilliQ water 3X and mounted with Mowiol mounting media. Samples were imaged with 3i CSU-W1 spinning disk microscope equipped with a Hamamatsu sCMOS Orca Flash4.0 camera using 60X /1.4NA plan-apochromat oil immersion objective and Abberior STED microscope equipped with a Photo Multiplier Tube using 100x/1.4NA Olympus UPLSAPO oil immersion objective.

### Live imaging

For investigating the FA turnover, WT and Vim^−/−^ fibroblasts were transfected with emerald-paxillin and allowed to reach confluency on a 22 mm coverslip. Upon reaching confluency and after 48 hrs of transfection, the coverslip was transferred to Atto Fluor (Thermo Fischer Scientific) live cell imaging chamber. Complete DMEM growth media was added and a scratch was made using a 10ul pipette tip. This media was replaced with live imaging media (1ml) from Invitrogen supplemented with L-glutamine, Pencillin, Streptamycin and 10% FBS and kept in the incubator for about 1h to allow the cells to begin migration. TIRF live imaging was performed on DeltaVision OMX with a Ring-TIRF system equipped with 3x Front illuminated sCMOS using 60x/1.49 NA Olympus APO N TIRF objective using 488 laser excitation. Imaging was done for 5-8 cells per imaging sessions of 1 hr with an interval of 20 secs, exposure time of 100-150ms, laser power at 1%, frequency of 96.5 Mhz, and image size of 1024×1024 pixels.

For investigating the FA interaction with vimentin, Vim^−/−^ fibroblasts were transfected with emerald-paxillin/GFP-FAK and mcherry-vimentin. Vim^−/−^ fibroblasts were chosen instead of WT fibroblasts to avoid vimentin overexpression artifacts. TIRF imaging was performed for 15 min with an interval of 3-5 secs, exposure time of 100-150ms, laser power at 1%, frequency of 96.5 Mhz, and image size of 1024×1024 pixels. Excitation wavelengths of 488nm and 568 nm were used for emerald-paxillin/GFP-FAK and mcherry-vimentin, respectively. For confocal live imaging, a Zeiss LSM880 equipped with 32-channel Airyscan Hexagonal element using 40x/1.2 NA Zeiss LD LCI Plan-Apochromat water immersion objective using 488 and 561 laser lines for Emerald-paxillin and mcherry-vimentin, respectively. The samples were imaged for 1h at 1 min intervals with optimal z spacing of 0.18 microns. All live imaging was performed in a temperature-controlled chamber at 37 C and 5% CO2, equipped with a humidifier.

For measuring the turnover of FA components, Emerald-paxillin transfected WT and Vim^−/−^cells were imaged with Zeiss LSM880 scope with 40x/1.2 NA Zeiss LD LCI Plan-Apochromat water immersion objective. Imaging was performed 24-48 h post transfection with 488nm laser line at 2% power. The settings including zoom and rotation were adjusted to include FAs, background, and cytoplasmic staining in each field of view and ensure that the time period of scanning was 100ms. Imaging was performed for 30 iterations with 2 pre-bleach and 28 post bleach scans, using rectangular areas on FAs for bleaching with 100% laser intensity. Intensities of bleached, background and reference regions over time were measured in ImageJ. Cytoplasmic region devoid of any FAs were used as reference region. The analysis was performed using easyFRAP standalone software (Rapsomaniki et al., 2012). The data was fitted with a single exponential fit.

### iPALM

HDF fibroblasts were plated on 25mm diameter round coverslips containing gold-nanorod fiducial markers (Nanopartz, Inc.) [1,2,3], coated with a thin layer of SiO2. After fixation and staining, an 18 mm coverslip was placed on top of the sample in STORM-buffer [4]. Alexa Fluor 647 channel was excited with 3 kW/cm2 intensity 647 excitation laser at an exposure of 30 ms and 100,000 were collected. Similarly, 561 channel stained with CF583R dye-labelled antibody was illuminated with 1–2 kW/cm2 intensity 561 excitation laser line for 50 ms exposure time and 100,000 frames were collected. Photo-switching of CF583R dye was induced using 2–10 W/cm2 intensity 405 laser excitation.

The collected frames were drift corrected and processed using PeakSelector software (Janelia Research Campus). For further analysis, the images were exported in ASCII format using PeakSelector software and analyzed using a custom Python script. Python script is available in this GitHub repository: https://github.com/eletca/iPALM-data-processing.

### Image Analysis and Statistics

Fixed cell images were analyzed using ImageJ (Collins, 2007; Schindelin et al., 2012) and custom-made macros. TIRF Videos were analyzed using FA Analysis Server (FAAS) (Berginski & Gomez, 2013). iPALM images were analyzed and visualized using a custom written Python program. Code is available in GitHub (https://github.com/eletca/iPALM-data-processing). GraphPad Prism was used for statistical analysis. Statistical analysis was performed using Welch’s t-test and Wilcoxon rank-sum test. The 3D iPALM data in the ASCII file was adjusted for the Z zero position using gold beads as fiducial markers. The zero level on the glass coverslip was set as the average Z position of the gold bead centers minus twice the standard deviation, as this serves as an approximation for the size of the gold beads. After performed correction, GB were removed not to interfere further analysis. Total 58 adhesions were selected manually from 3 different cells and histograms were plotted for each adhesion using a custom python script to define Z centers of vimentin and vinculin distribution. Taking into account the sample preparation, gold beads were covered approximately by 100nm SiO2 layer, final Z center values should be considered approximately 100nm less than the displayed results due to uncertainty of the glass layer thickness distribution.

To perform an unbiased classification of focal adhesion (FA) subpopulations, time-lapse Videos of Vim^−/−^ MEFs co-expressing mCherry-vimentin and GFP-FAK were analyzed. Individual FA tracks were first identified and quantified using a custom Python script available here (Github). The resulting track data from 10 cells were pooled, and tracks with a lifetime of less than 20 frames (100 seconds) were excluded from the analysis to ensure only persistent FAs were considered, yielding a final dataset of 481 high-confidence tracks. For each track, another custom Python script was used to calculate eight distinct features summarizing its dynamic profile. These features were: (1) average area (µm²), (2) average velocity (µm/s), (3) directional persistence, (4) directionality, (5) average mean FAK intensity normalized to the first frame, (6) average mean vimentin intensity normalized to the first frame, (7) average Pearson’s Correlation Coefficient (PCC), and (8) average Intensity Correlation Quotient (ICQ).

Time-lapse videos were analyzed using dense optical flow (Farnebäck method) between consecutive grayscale frames after downsampling to a fixed width for computational stability. For each frame pair, mean flow magnitude (px/frame) was used as a proxy for overall vimentin network motion and converted to px/s using the acquisition frame rate. Contraction-like behavior was quantified by computing the spatial divergence of the flow field (∂fx/∂x + ∂fy/∂y), where negative values indicate net local convergence; results were summarized as mean divergence and the fraction of frame transitions with negative divergence (“contraction fraction”). Coiling or torsional motion was assessed using a vorticity proxy, calculated as the absolute curl magnitude |∂fy/∂x − ∂fx/∂y| and averaged over the field, with higher values reflecting stronger rotational components. Directional heterogeneity was quantified by circular variance of flow angles computed on the top 30% most mobile pixels, where low and high values indicate aligned versus multidirectional motion, respectively. High-motion outliers (“bursts”) were defined as frame transitions exceeding mean + 2 standard deviations of flow magnitude, capturing transient, high-amplitude rearrangement events.

Prior to clustering, any missing feature values were imputed using the median for that feature, and the entire feature set was standardized using a Z-score transformation (mean of 0, standard deviation of 1) to ensure all features contributed equally to the analysis. Unsupervised classification was performed using a Gaussian Mixture Model (GMM) with three components, a choice supported by the Bayesian Information Criterion (BIC). The resulting cluster assignments were then used to analyze the distinct phenotypic properties of each FA subpopulation. To formally test for significance between the identified clusters, a Kruskal-Wallis test was performed for each dynamic feature, followed by Dunn’s post-hoc test for pairwise comparisons. The Kruskal-Wallis test, a non-parametric equivalent of ANOVA, was chosen as it does not assume a normal (Gaussian) distribution of the data. Many types of biological measurements from microscopy, such as area or intensity, are often skewed and do not follow a perfect bell-shaped distribution, making this rank-based test a more robust and scientifically safer choice. The most prototypical examples for each cluster were identified by calculating the Euclidean distance of each track from its assigned cluster’s centroid in the standardized feature space.

### Live-cell imaging and quantitative analysis of vimentin–focal adhesion dynamics

Live-cell imaging of fibroblasts co-expressing vimentin–mCherry and paxillin–GFP was performed using spinning-disk confocal microscopy at 37 °C and 5% CO₂. Image sequences were acquired at 10–15 s intervals for 15–20 min using a 100×/1.4 NA oil-immersion objective.

Quantitative analysis of vimentin–focal adhesion (FA) interactions was performed on the time-lapse datasets using ImageJ/Fiji and custom Python scripts. For each frame, Pearson’s correlation coefficients were calculated between vimentin (magenta) and FA (green) channels to quantify temporal changes in colocalization. Centroid positions of individual adhesions and nearby vimentin filament segments were determined from thresholded masks, and frame-to-frame distances between centroids were computed to assess dynamic proximity.

Intensity profiles for both channels were extracted from identical regions of interest and plotted as a function of time to evaluate synchronous or asynchronous signal fluctuations. Correlation and intensity traces were analyzed to identify periodic patterns of association and dissociation, corresponding to adhesion assembly–disassembly cycles.

Data from at least three independent recordings were analyzed. All image processing and quantitative analyses were conducted using identical thresholding and filtering parameters.

The plots of Pearson correlation and centroid distance distributions were generated using custom Python scripts (available upon request) to visualize temporal fluctuations and spatial proximity between vimentin and FA structures (Fig. S5).

### Mass spectrometry and Integrative interactome analysis

For co-immunoprecipitation and mass spectrometry, neonatal HDFs (PCS-201-010, ATCC) cells were cultured in 15 cm dishes in Dulbecco’s modified media (DMEM, Sigma #D6171) supplemented with 10 % fetal bovine serum (FBS, Biowest #S1810), 2 mM of L-glutamine (Biowest #X0550), 100 U/ml of penicillin, 100 µg/ml of streptomycin (Sigma #P0781) at 37 °C in a 5% CO2 incubator until approximately 80% confluence. Cells were then starved for 1 h in RPMI medium lacking glucose and amino acids (Amersham). Subsequently, cells were treated for 30 minutes with a mixture containing 1 mM L-glutamine (Sigma), 2.5 mM glucose (Gibco), 1× essential amino acids (Gibco), and 100 nM insulin (Sigma). Cells were washed twice with PBS and lysed in buffer containing 0.5% NP-40, 150 mM NaCl, 20 mM HEPES (pH 7.4), and 1 mM EDTA supplemented with protease and phosphatase inhibitors (Thermo). Lysis was performed for 1 h at 4 °C with rotation. Lysates were cleared by centrifugation at 14,000 rpm for 10 minutes, and the supernatants were transferred to fresh tubes. Protein concentrations were determined using a BCA assay kit (Thermo). Protein G beads (Thermo) were washed twice and equilibrated in lysis buffer at a 1:1 ratio. For pre-clearing, lysates were incubated with 7 µl of beads for 1 h at 4 °C with rotation, after which the beads were removed. Antibody-coupled beads were prepared separately by washing protein G beads four times with PBS and incubating them with either V9 antibody or mouse IgG control (8 µg antibody per 1 mg beads) in PBS for 1 h at 4 °C with rotation. Following incubation, beads were washed and resuspended in lysis buffer at a 1:1 ratio. Pre-cleared lysates were then incubated with 25 µl of antibody-conjugated beads for 4 h at 4 °C with rotation. After binding, beads were sequentially washed twice with lysis buffer, twice with PBS, and twice with Milli-Q water. Proteins bound to the beads were digested on-bead by adding trypsin (5 µg/ml) in six volumes of 2 M urea prepared in 25 mM ammonium bicarbonate (pH 7.5) and incubating for 30 min at 27 °C. The supernatant was collected, and the beads were washed twice with 2.5 volumes of 2 M urea in 25 mM ammonium bicarbonate containing 1 mM DTT. All supernatants were combined and allowed to digest overnight. The following day, cysteine residues were alkylated by adding 20 µl iodoacetamide (5 mg/ml) and incubating for 30 minutes at room temperature in the dark. Peptides were then acidified with TFA and desalted using a 96-well plate (Waters) according to the manufacturer’s instructions. The samples were dried on a speed-vac centrifuge and kept at –80 °C. Prior to analysis, the samples were dissolved in 0.1 % formic acid. Approximately 100 µg of each sample was loaded for LC-MS/MS analysis. The analysis was conducted using an Easy-nLC 1000 liquid chromatograph (Thermo Scientific) coupled to an Orbitrap Fusion Lumos Tribrid Mass Spectrometer (Thermo Scientific). The peptides were loaded on a pre-column (0.1 × 20 mm), followed by separation in an analytical column (75 μm x 150 mm), both packed with 5 μm Magic C18 silica particles (Michrom). The peptides were separated using a 60-minute gradient (5-42 % B in 65 min, 42-100 % B in 5 min, 100% B for 3 minutes) in which solvent B was acetonitrile/water (95:5), at a flow rate of 300 nl/min.

MS/MS data were acquired in positive ionization mode using data-dependent acquisition. The MS survey scans were acquired with a resolution of 70 000 across the range of 300–2000 m/z, an AGC target of 106 and a maximum fill time of 120 ms. Up to 10 of the most intense ions with charge >+1 were selected for HCD fragmentation using isolation window of 2.0 m/z and with AGC target of 5 × 104 and a maximum fill time of 240 ms. MS/MS spectra were recorded with 17 500 resolution, and dynamic exclusion window of 20 s was used. The raw mass spectrometry data have been deposited to the ProteomeXchange Consortium via the PRIDE29 partner repository (Perez-Riverol et al., 2025) with the dataset identifier PXD075418. The AP-MS dataset can be accessed through the PRIDE repository using the following reviewer credentials (**Username; Password: JjIzPhxawywV**). The dataset will be made publicly available upon manuscript acceptance.

Raw files obtained from the LC-MS/MS analyses were processed using MaxQuant software (version 2.0.1.0) (Cox & Mann, 2008; Tyanova et al., 2016a) and searched against a SwissProt human protein database with added common contaminants using the built-in Andromeda search engine (Cox et al., 2011). Trypsin digestion with a maximum of 2 missed cleavages, cysteine carbamidomethylation as a fixed modification, and methionine oxidation as variable modification were selected as the parameters of these searches. “Match between runs” option in MaxQuant was used with matching time window of 2 minutes. and alignment time window of 20 minutes. The Peptide level false discovery rate (FDR) was set to 1 % and was determined by searching against a concatenated normal and reversed sequence database. Label-free quantitation was performed using the fast LFQ algorithm (Cox et al., 2014). Otherwise, the default settings in MaxQuant were used in data processing. The database search results with LFQ intensities were analysed using Perseus (1.6.7.0)(Tyanova et al., 2016b). Proteins only identified by site, proteins considered contaminants and reverse were discarded. Only proteins with 3 values in at least one group (stv or nut) were kept. Data was transformed and proteins with a negative-control LFQ/(average vimentin-IP LFQ) ratio >1 were removed. Interactors were scored with MIST(Hu et al., 2017) and gene ontology analysis was conducted with Panther(Mi et al., 2019). Vimentin-associated proteins were identified from AP-MS datasets (MiST ≥ 0.9). Only interactors previously linked to focal adhesion or actin-regulatory functions were retained for network visualization (Fig. S9; Table 1).

## Acknowledgments

We thank Prof. Johanna Ivaska for valuable comments on the manuscript and Prof. Lea Sistonen for practical support. We acknowledge the invaluable assistance provided by the team of the Advanced Imaging Center, Janelia Research Campus to use iPALM microscope and further image processing, with special thanks to AIC Director Teng-Leong Chew. This study was supported by Academy of Finland, Sigrid Jusélius Foundation, Magnus Ehrnrooth Foundation, the Endowment of the Åbo Akademi University, K. Albin Johanssons stiftelse, Maud Kuistila Memorial Foundation, Liv och Hälsa Foundation, and the Foundation “Konung Gustaf V:s och Drottning Victorias Frimurarestiftelse”. We also like to thank Cell Imaging and Cytometry Core, Turku Bioscience Centre, which is supported by the Finnish Advanced Microscopy Node of Euro-BioImaging Finland (Turku, Finland), Turku Bioimaging and Euro-Bioimaging Node at Turku Bioscience Center, Turku, Finland, and Biocenter Finland for the research facilities.

This work was supported by the Research Council of Finland, FIRI 2023 grant (decision numbers: 359073, 358879) and FIRI 2024 (grant decision numbers: 367582 and 367577).

Additionally, this research received support from the InFLAMES Flagships Programme of the Research Council of Finland (decision numbers: 337530, 337531, 357910, and 35791).

G.J was supported by grants awarded by the Academy of Finland (338537, 371287, and 374180), the Sigrid Juselius Foundation, the Cancer Society of Finland and Åbo Akademi University Research Foundation (G.J., CoE CellMech) and by Drug Discovery and Diagnostics strategic funding to Åbo Akademi University (G.J.).

## Author contributions

Experimental design was prepared by Arun P Venu and John E Eriksson. Scratch wound experiments and image analysis were performed by Arun P Venu and Joakim Edman. TIRF live imaging experiments were performed by Arun P Venu, Mayank Modi, and related data analysis were performed by Arun P Venu and Mayank Modi. Spinning Disc imaging and related image analysis were performed by Mayank Modi. Peiru Yang performed the SDS PAGE and western blots. STED imaging was performed by Arun P Venu with assistance from Elena Tcarenkova and Ujjwal Aryal. Giulia Sultana, Jenny Pessa, Mayank Modi prepared samples, conducted iPALM imaging and image processing. Arun P Venu and Elena Tcarenkova performed live cell and iPALM image analysis using Python script. Alexander Minin provided the rat fibroblasts and some of the data with those fibroblasts. Advice on how to examine the interactomics data was given by Leila Rato. Advice and assistance were provided by Guillaume Jacquemet and Fang Cheng. The manuscript was written by Arun P Venu, Mayank Modi, and John E Eriksson. All authors revised and edited the manuscript. John E Eriksson acquired the funding.

## Video Legends

**Video 1A.**
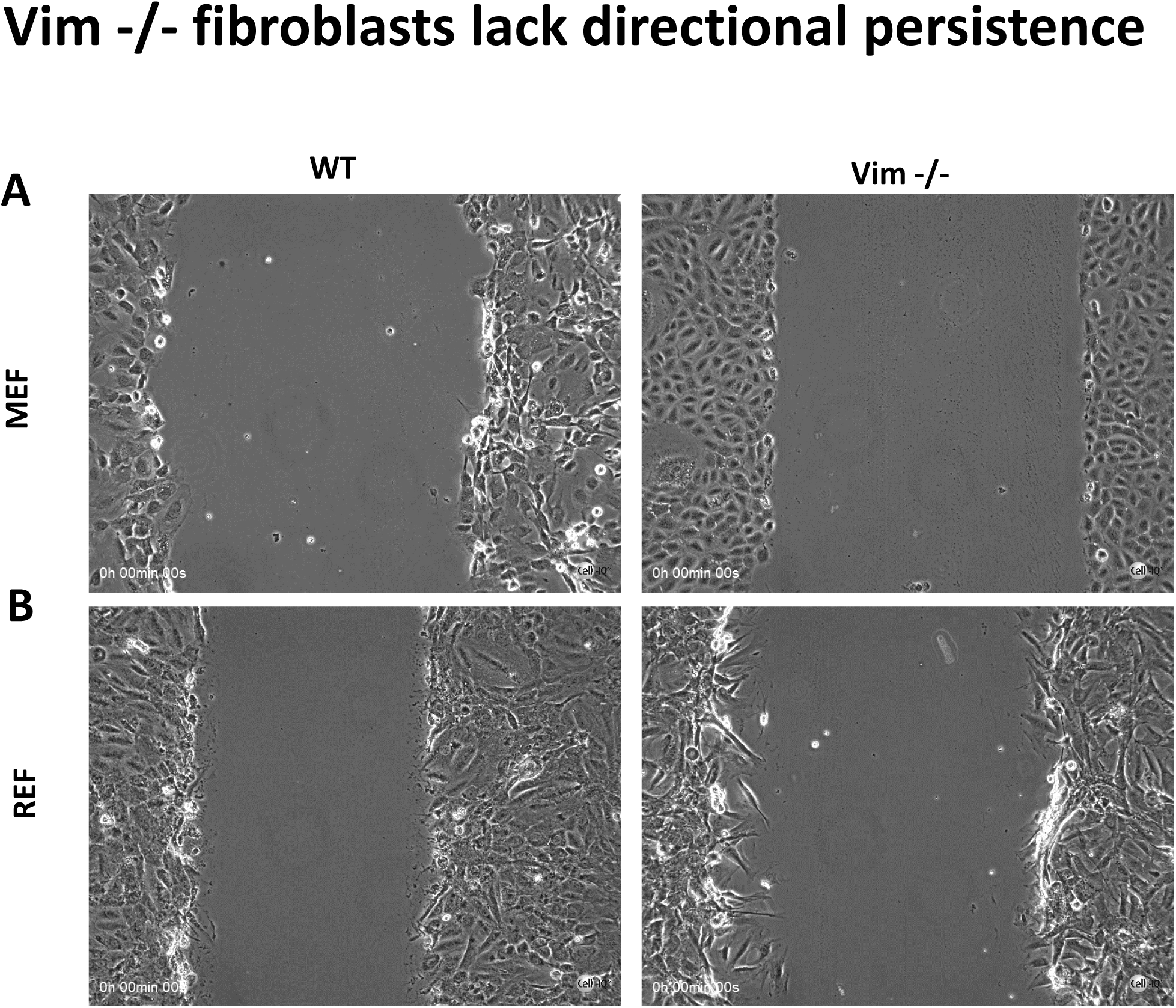
Scratch wound migration of WT and Vim^−/−^ MEF cells. **Video 1B**. Scratch wound migration of WT and Vim^−/−^ REF cells.

**Video 2.**
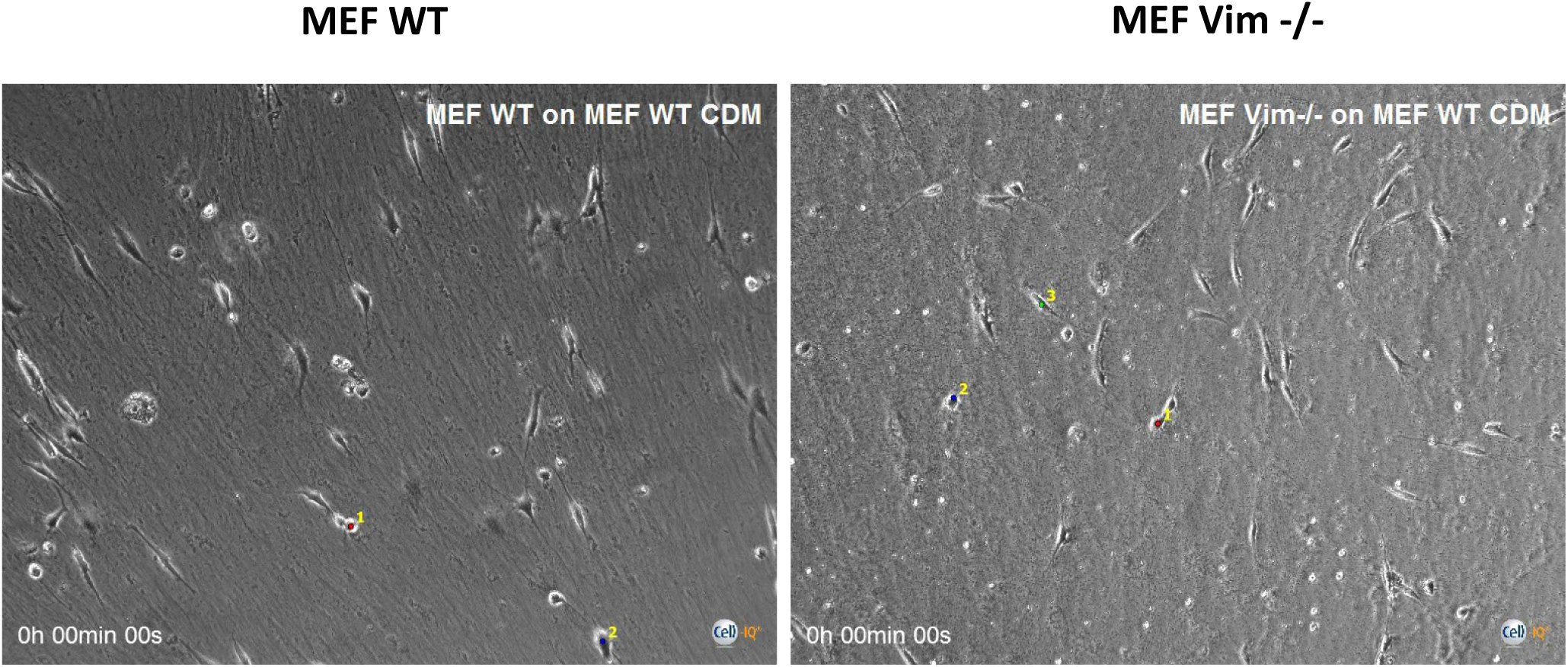
Migration of WT and Vim^−/−^ MEFs on WT cell-derived matrix (related to Fig. 1). Vim^−/−^ cells fail to align with CDM fibers.

**Video 3.**
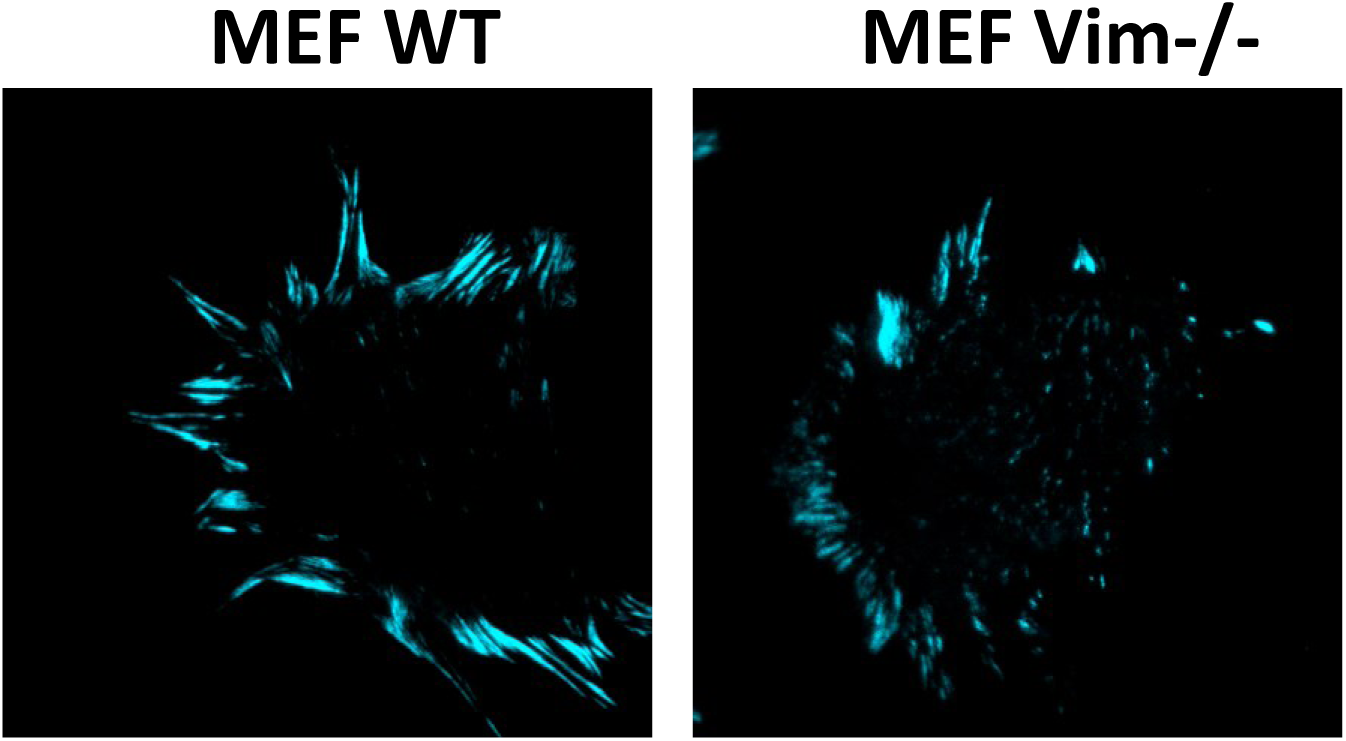
FA dynamics during directional migration (related to Fig. 2). Vim^−/−^ MEFs expressing Emerald-paxillin(pseudo-colored in cyan) show rapid and disorganized FA disassembly.

**Video 4A_4B.**
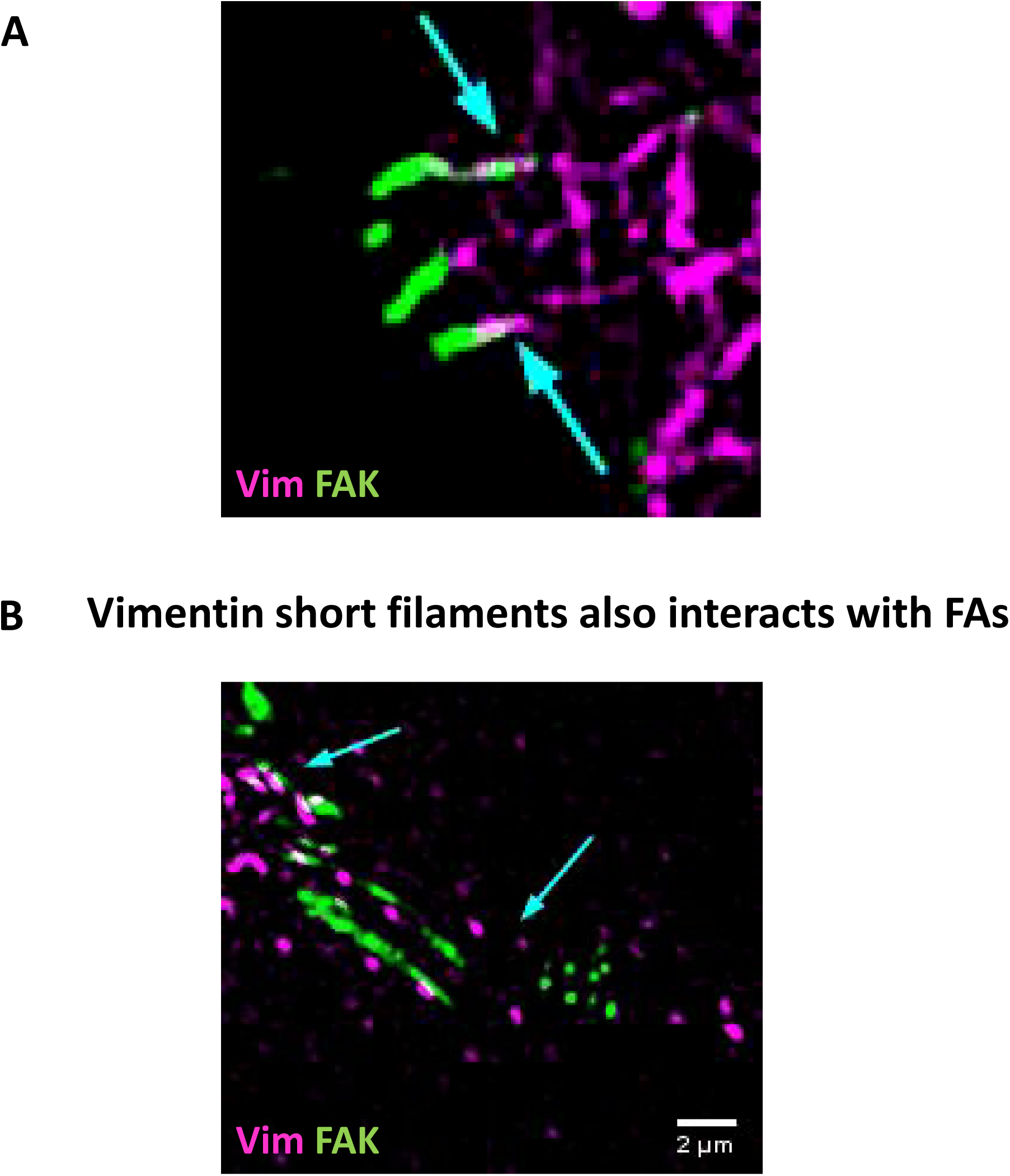
Dynamic interaction of vimentin filaments with maturing and retracting FAs in Vim^−/−^ MEFs co-expressing GFP-FAK and mCherry-vimentin(pseudo-colored in magenta) (related to Fig. 4).

**Video 5A_5B.**
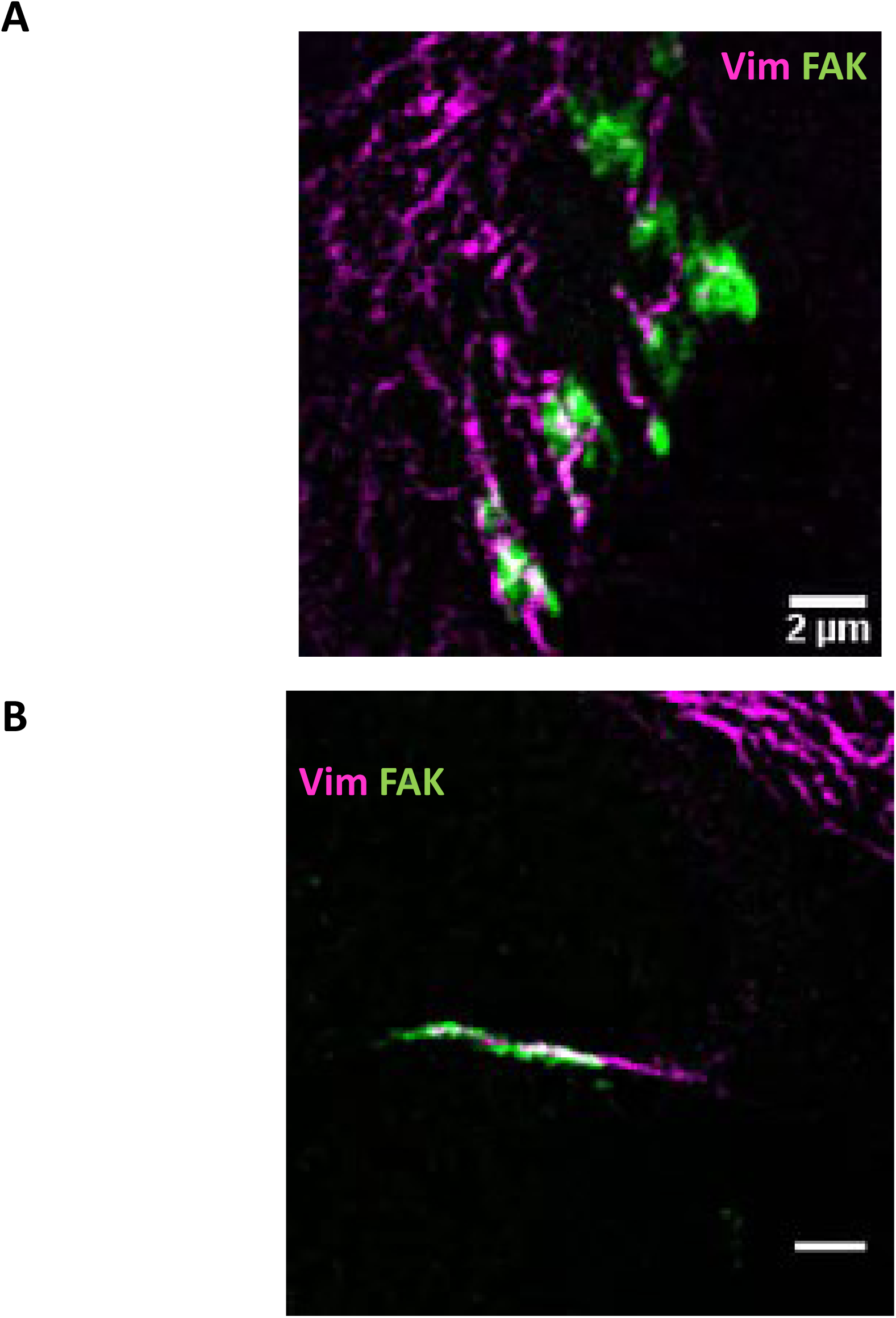
Dynamic interaction of vimentin filaments with maturing and retracting FAs in Vim^−/−^ MEFs co-expressing GFP-FAK and mCherry-vimentin(pseudo-colored in magenta) (related to Fig. 4).

**Video 6.**
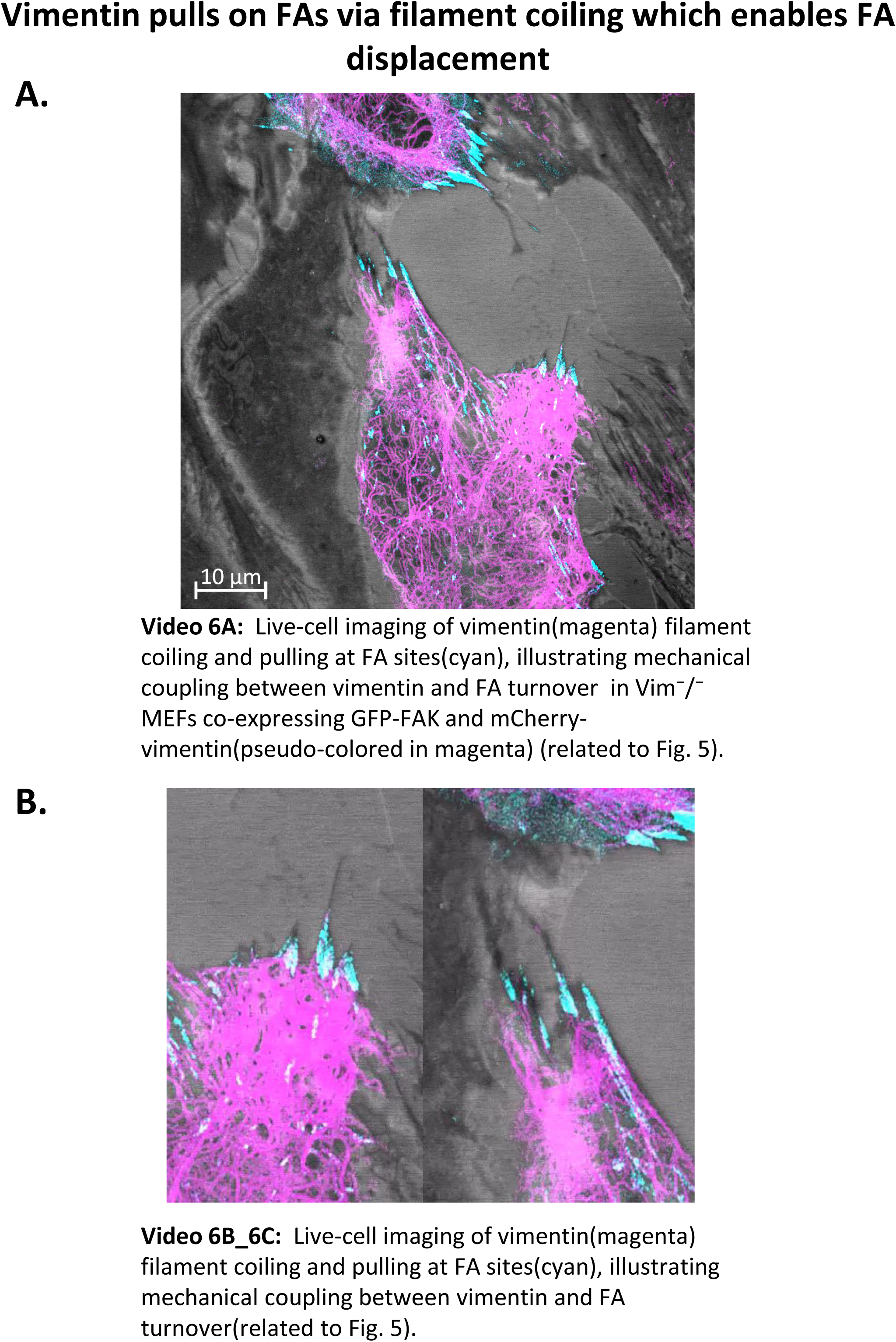
Live-cell imaging of vimentin filament coiling and retraction at FA sites, illustrating mechanical coupling between vimentin and FA turnover in Vim^−/−^ MEFs co-expressing GFP-FAK(pseudo-colored in cyan) and mCherry-vimentin(pseudo-colored in magenta) (related to Fig. 5).

**Video 7.**
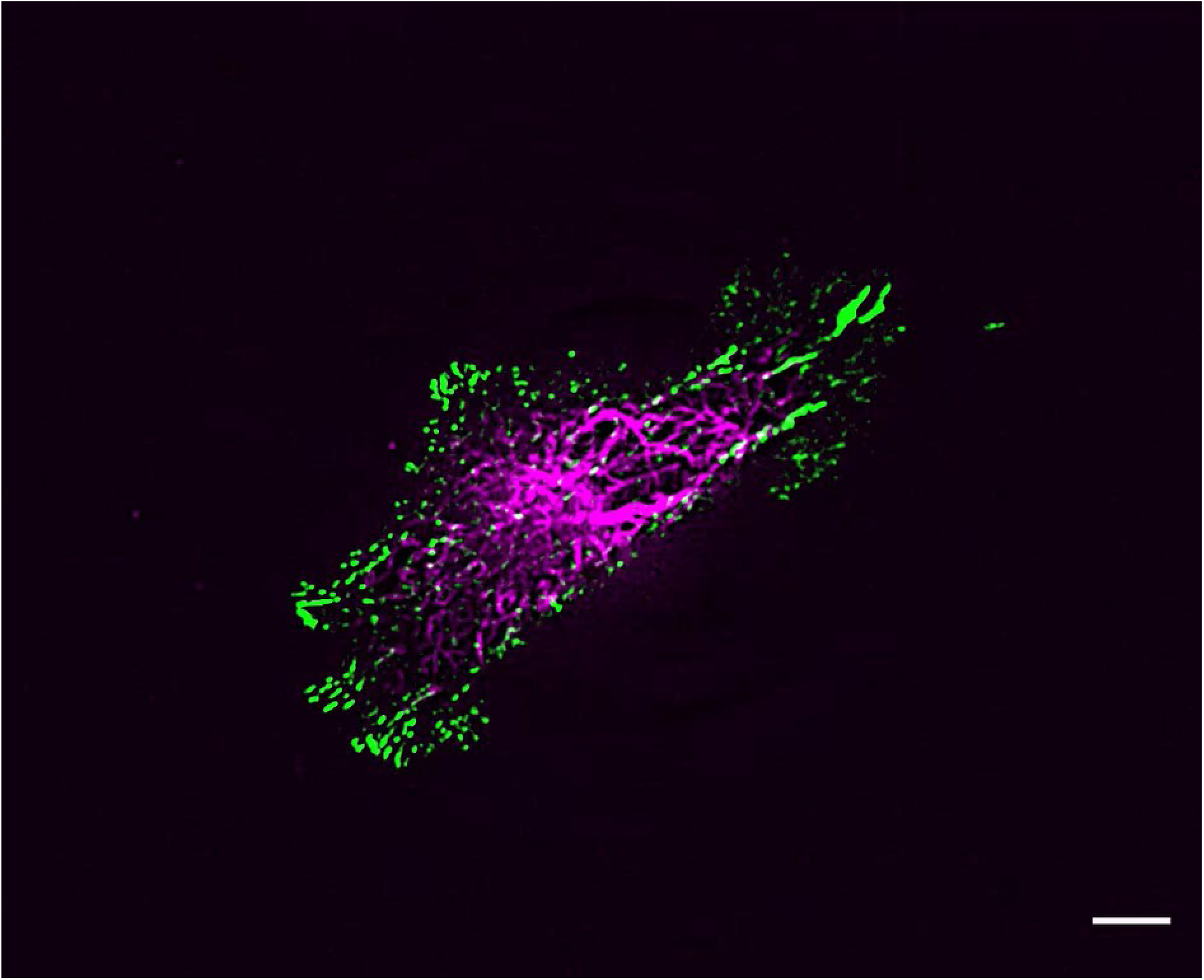
Live-cell imaging of vimentin filament dynamics at FA sites, illustrating mechanical coupling between vimentin and FA turnover in Vim^−/−^ MEFs co-expressing GFP-FAK and mCherry-vimentin(pseudo-colored in magenta) (related to Fig. 6).

**Video 8.**
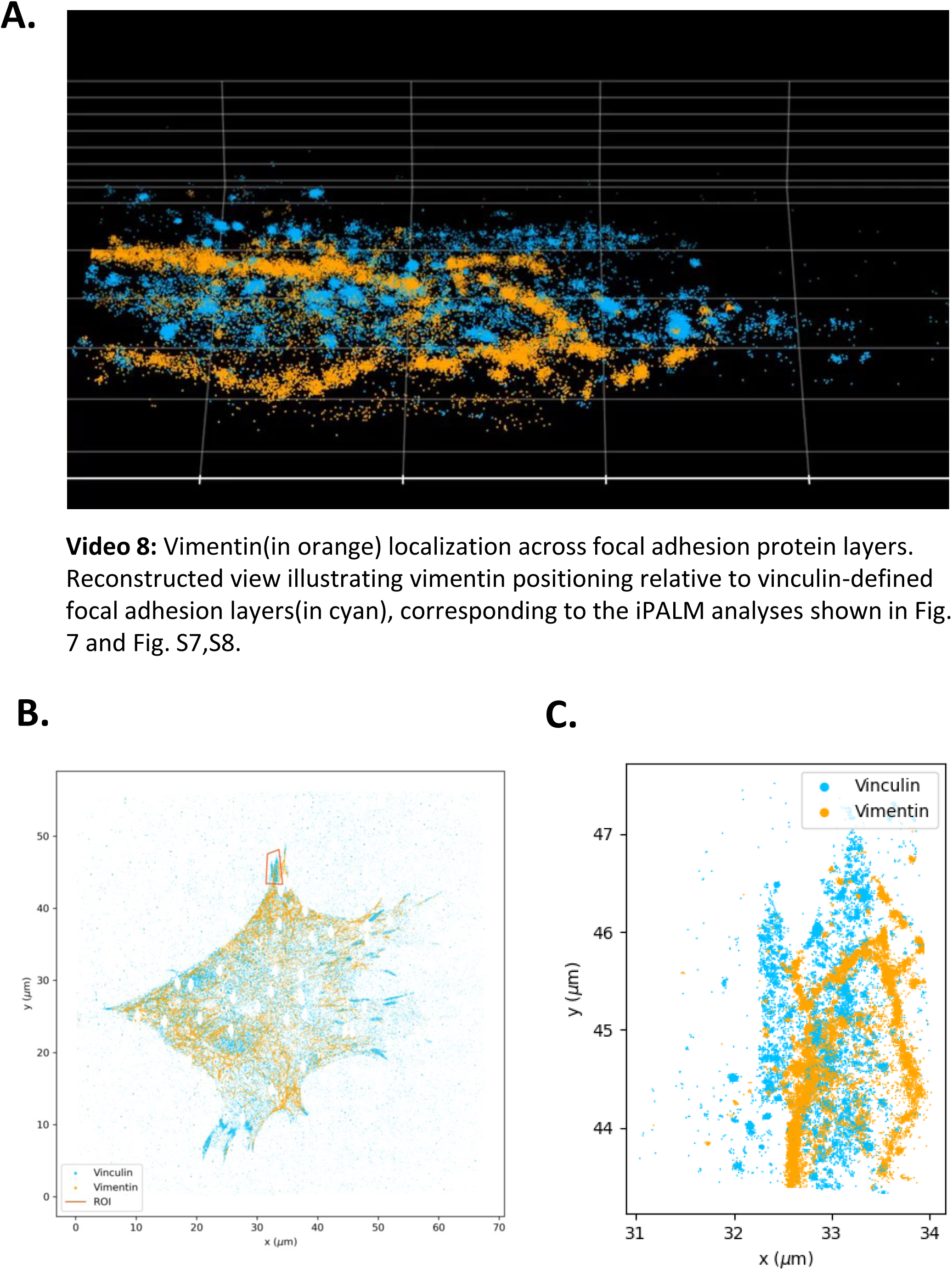
Vimentin localization across focal adhesion protein layers (related to Fig. 7 and Fig. S7, 8). Reconstructed view illustrating vimentin(in orange) positioning relative to vinculin-defined focal adhesion layers(in cyan), corresponding to the iPALM analyses shown in Fig. 7 and Fig. S7,8. Visualization highlights dynamic occupancy of vimentin across FA nanoarchitectural strata.

## Supplementary figures

**Figure S1.**
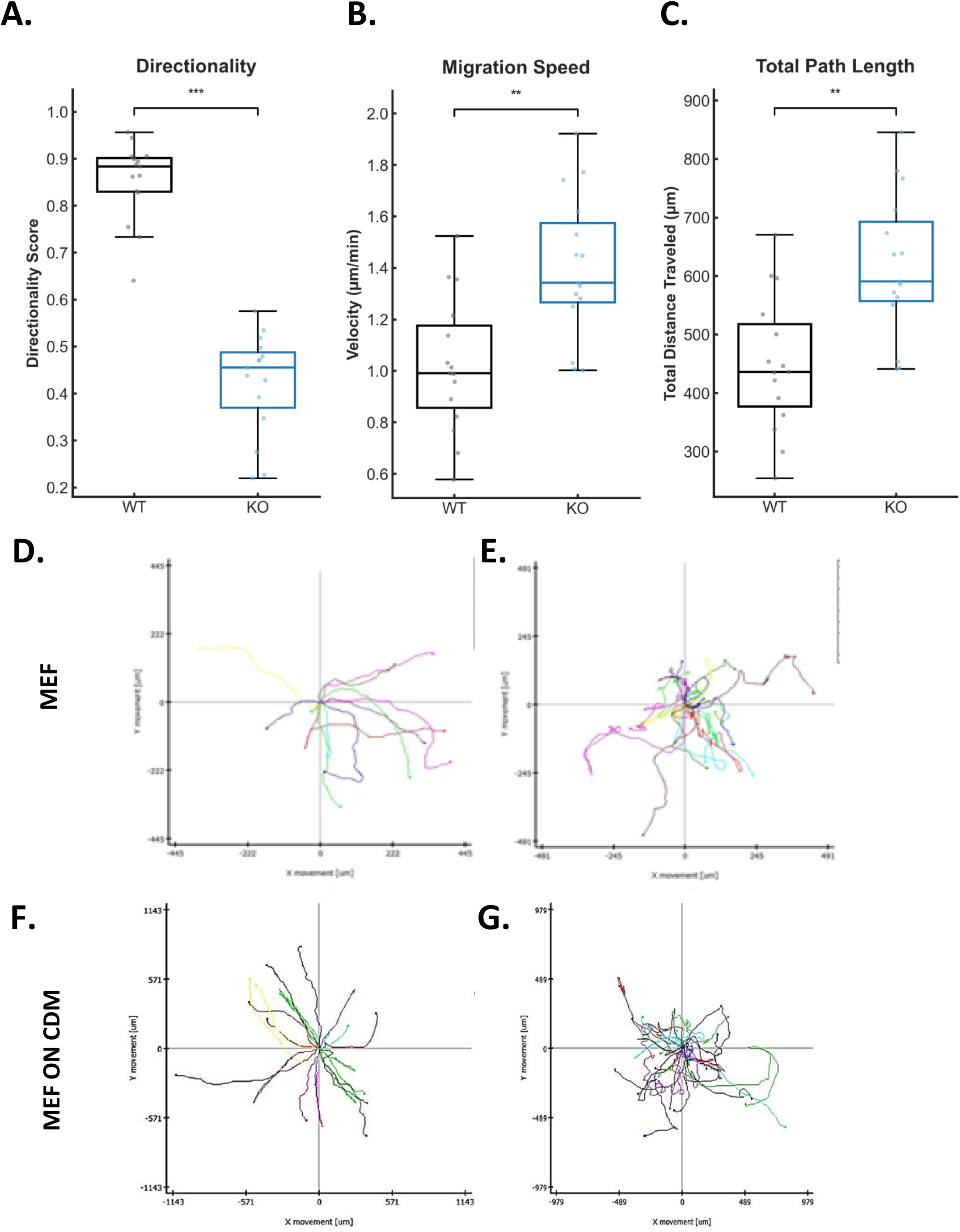
Directional migration defects in Vim^−/−^ MEFs during random migration and on cell-derived matrix. (A) Directionality score, (B) migration speed, and (C) total path length of WT and Vim^−/−^ MEFs during random migration. (D–E) Representative random migration trajectories of WT and Vim^−/−^ MEFs. (F–G) Migration trajectories of WT and Vim^−/−^ MEFs on WT cell-derived matrix (CDM). Vim^−/−^ cells exhibit reduced directionality despite increased speed and path length.

**Figure S2.**
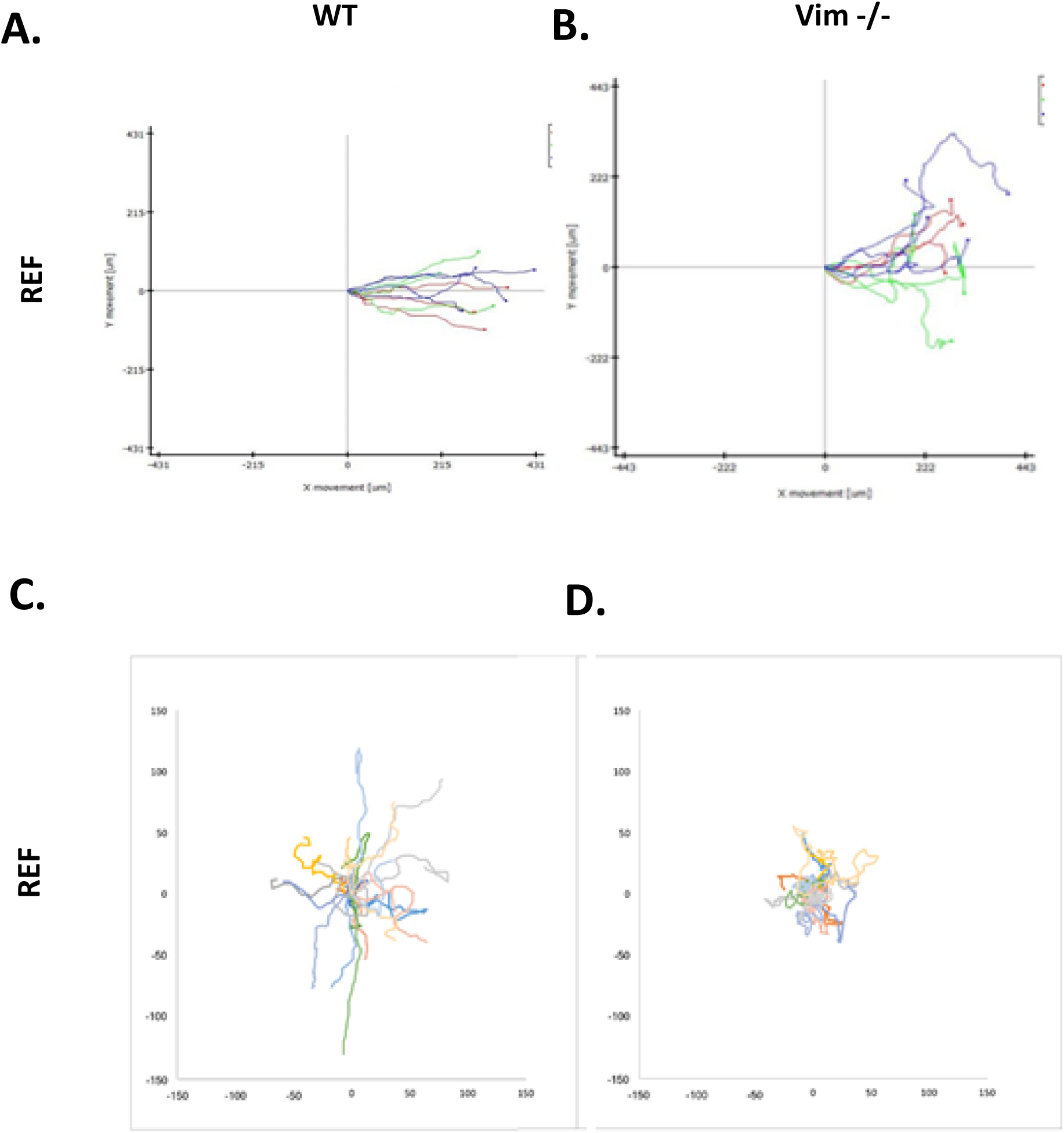
Vim^−/−^ rat embryonic fibroblasts (REFs) also lack directional persistence. (A–B) Aligned migration trajectories of WT and Vim^−/−^ REFs during wound-directed migration. (C–D) Random migration trajectories of WT and Vim^−/−^ REFs. Loss of directional persistence in Vim^−/−^ cells is conserved across species.

**Figure S3.**
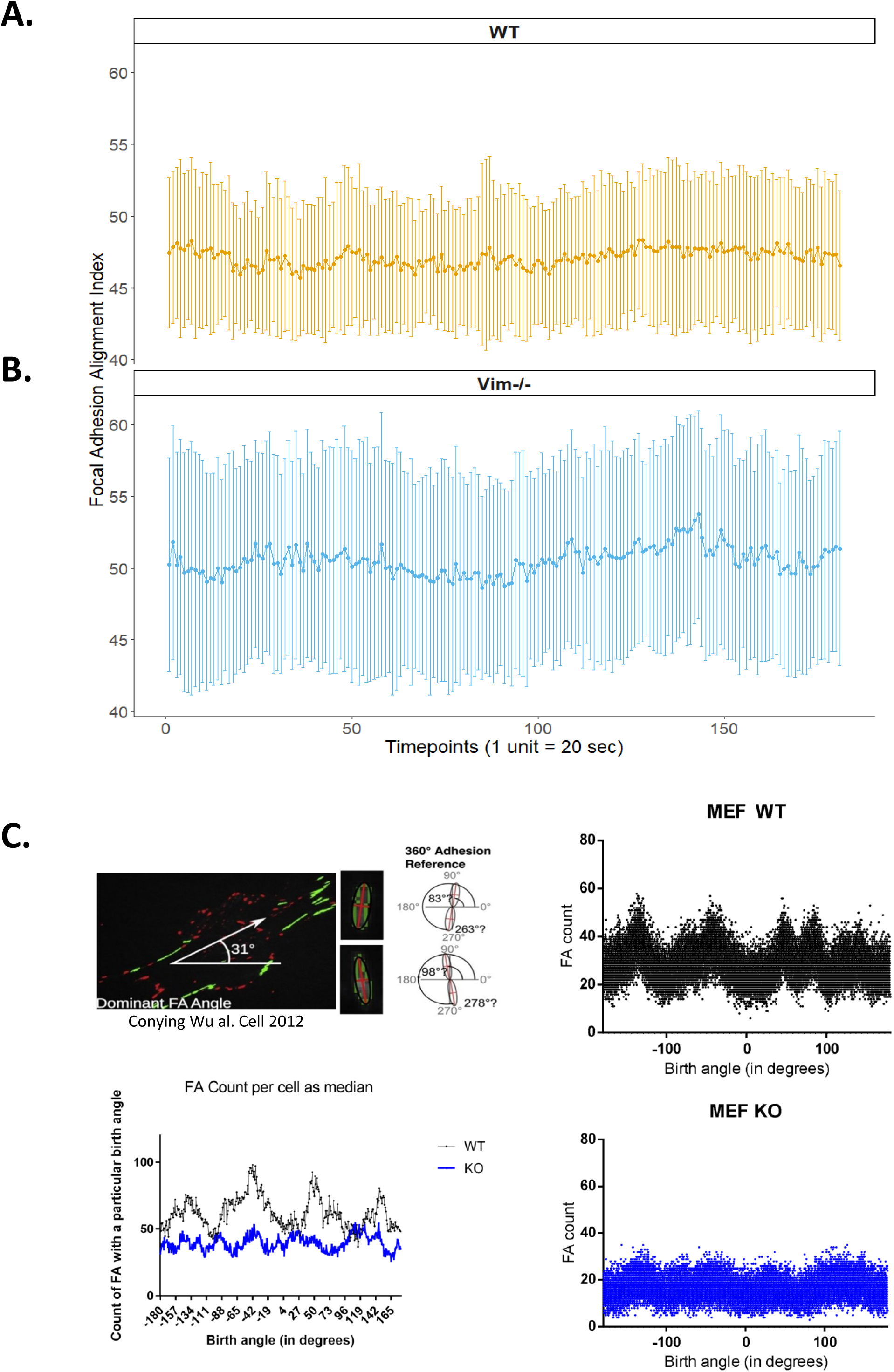
Temporal instability and loss of angular periodicity of focal adhesions in Vim^−/−^ MEFs. (A–B) Time-resolved FAAI traces for WT and Vim^−/−^ MEFs. Vim^−/−^ cells display increased temporal fluctuations. (C) Analysis of FA birth angles reveals periodic angular organization in WT cells but a near-uniform distribution in Vim^−/−^ cells.

**Figure S4.**
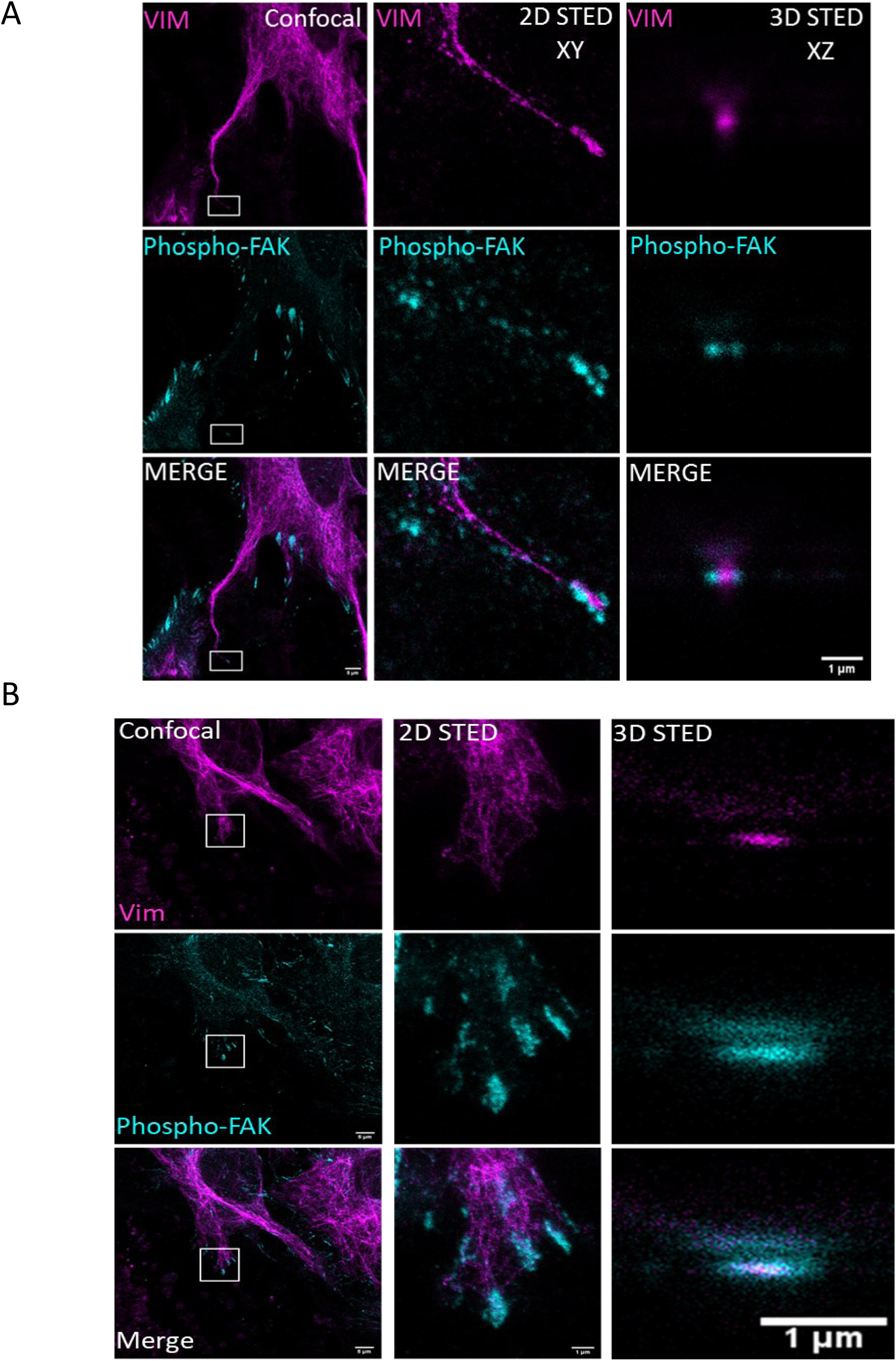
Vimentin associates with phosphorylated FAK during focal adhesion maturation and retraction. (A) Confocal and super-resolution STED imaging of Vim^−/−^ MEFs expressing vimentin (magenta) and stained for phospho-FAK (cyan). Insets indicate focal adhesions selected for high-resolution analysis. (B) Two-dimensional (XY) and three-dimensional (XZ) STED reconstructions reveal spatial proximity of vimentin filaments to phospho-FAK–positive adhesion sites during distinct stages of adhesion maturation and retraction. Scale bar, 1 µm.

**Figure S5.**
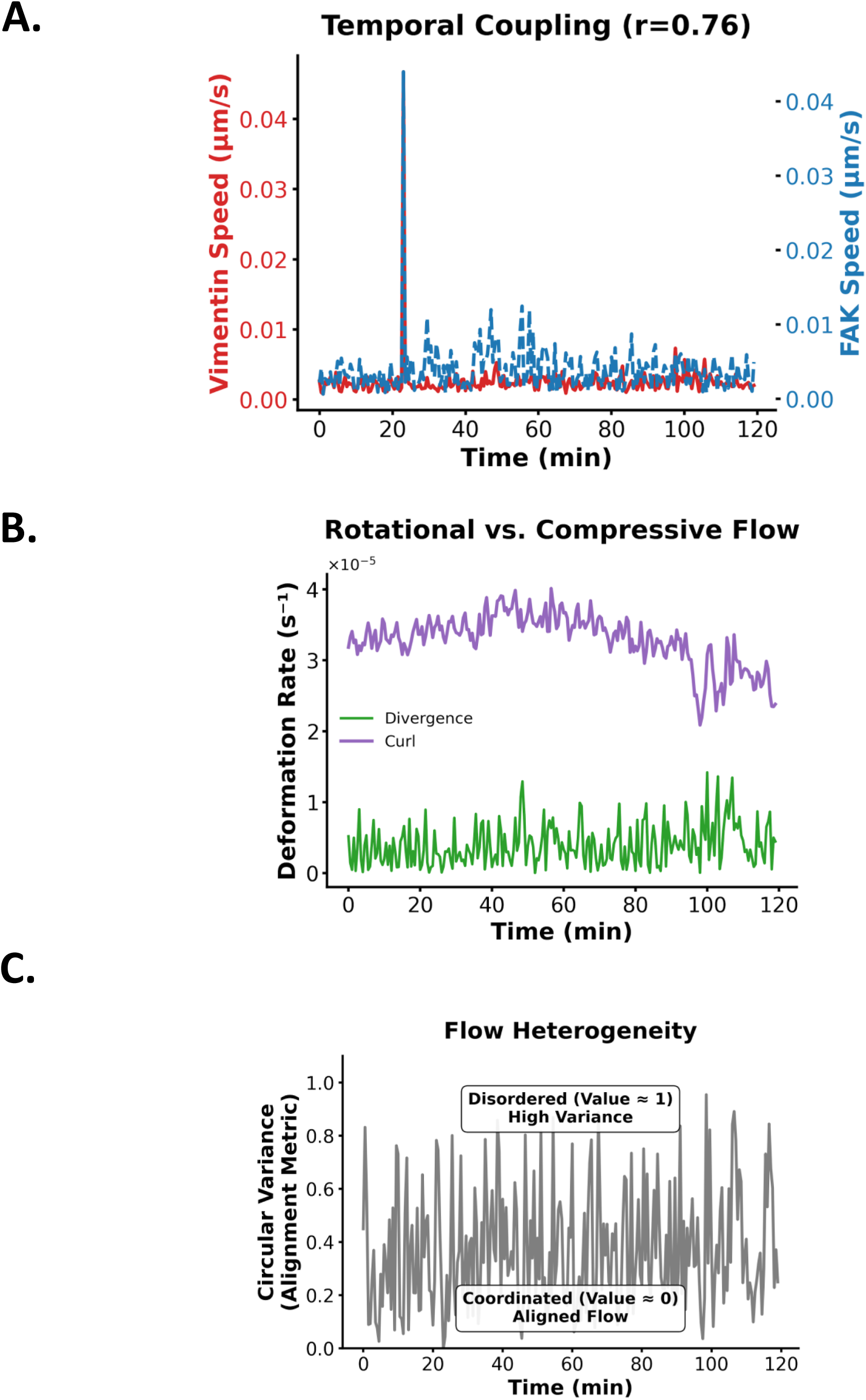
Vimentin pulls on focal adhesions via filament coiling to enable FA displacement. (A) Temporal Coupling of Cytoskeletal and Adhesion Dynamics. Dual-axis time-series plot overlaying vimentin remodeling speed (red, left axis) and FAK displacement speed (blue dashed, right axis). The peaks in vimentin activity coincide temporally with peaks in focal adhesion sliding, providing visual confirmation that vimentin remodeling is tightly coupled to adhesion dynamics in real-time. (B) Rotational vs. Compressive Flow Components. Decomposition of the flow field into Curl (Rotational component, purple) and Divergence (Isotropic contraction/expansion, green). The magnitude of rotational flow consistently exceeds compressive flow, indicating that cytoskeletal stress is managed primarily through local twisting and buckling of filaments rather than global shrinkage. Labels indicate the dominant deformation mode for each component. (C) Quantification of flow directionality using Circular Variance (0 = Aligned, 1 = Disordered). The network cycles between phases of highly coordinated flow (low variance, value ≈ 0) and disordered reorganization (high variance, value ≈ 1), suggesting that remodeling occurs in intermittent bursts of collective filament movement.

**Figure S6.**
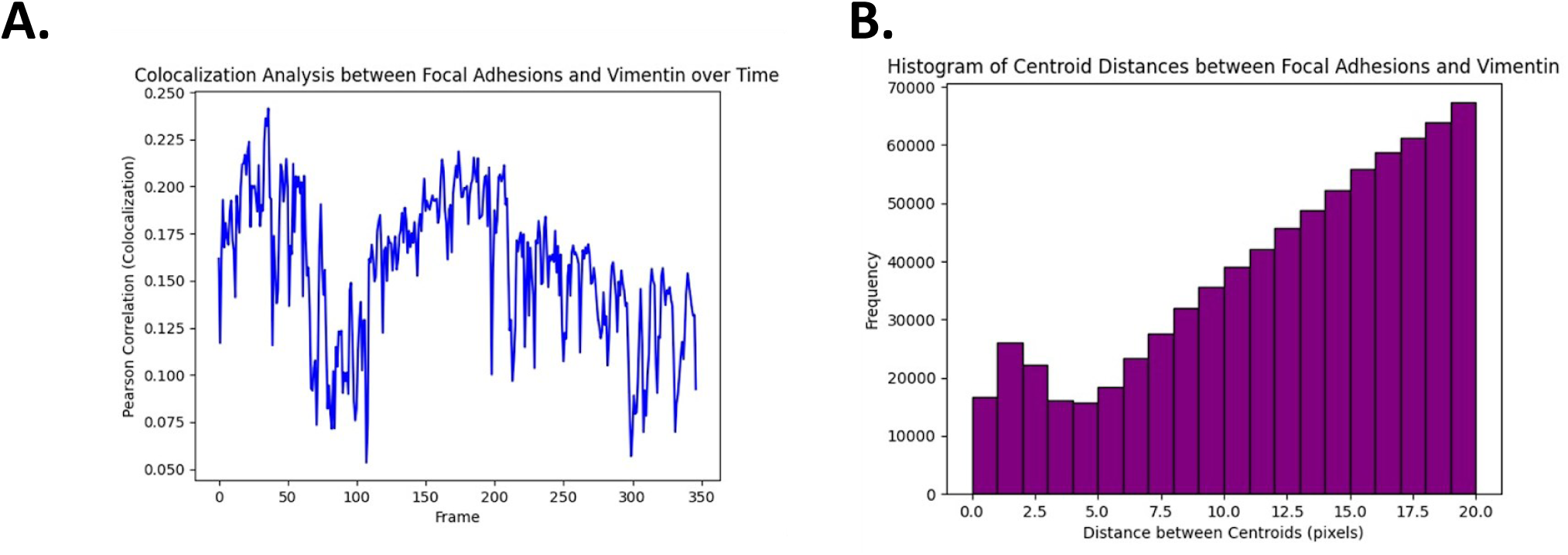
Time-resolved colocalization analysis of vimentin and focal adhesions during migration. (A) Time-dependent Pearson correlation coefficient (PCC) between vimentin and FA markers across individual adhesion lifetimes. (B) Corresponding intensity correlation quotient (ICQ) analysis reveals dynamic changes in vimentin-FA association over time, indicating transient and stage-specific coupling during FA maturation and turnover.

**Figure S7.**
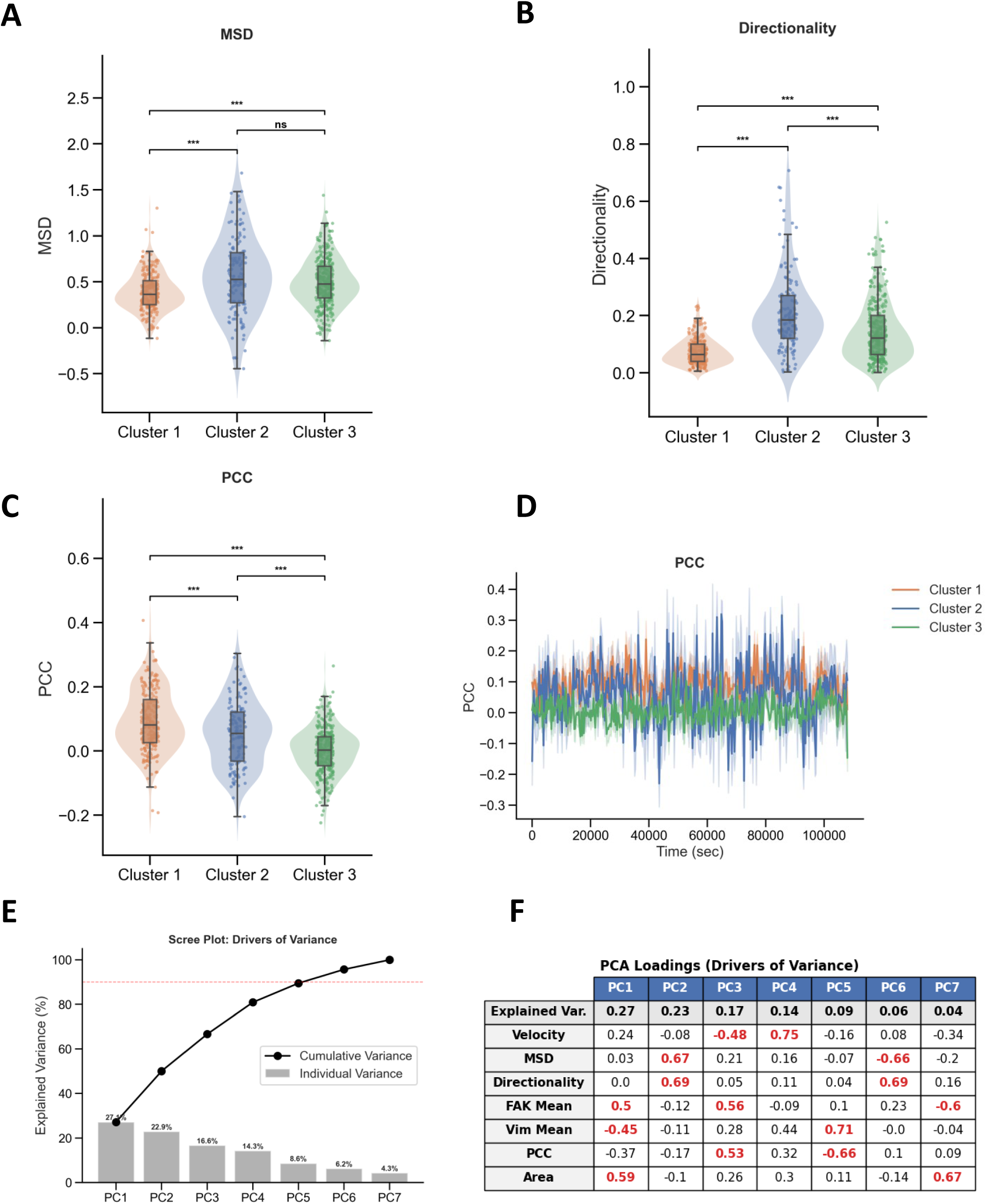
Biophysical properties and validation of adhesion clusters. (A-C) Raincloud plots quantifying (A) Mean Squared Displacement (MSD) $\alpha$-exponent, (B) Directionality ratio, and (C) Pearson Correlation Coefficient (PCC) between Vimentin and FAK. Cluster 1 (orange) exhib its the lowest MSD and Directionality (anchored) but the highest PCC (strong coupling). (D) Time-resolved analysis of the Vimentin-FAK Pearson Correlation Coefficient (PCC). (E) Scree plot showing the variance explained by each Principal Component. (F) PCA Loadings table identifying the specific features driving the separation along the principal axes.

**Figure S8.**
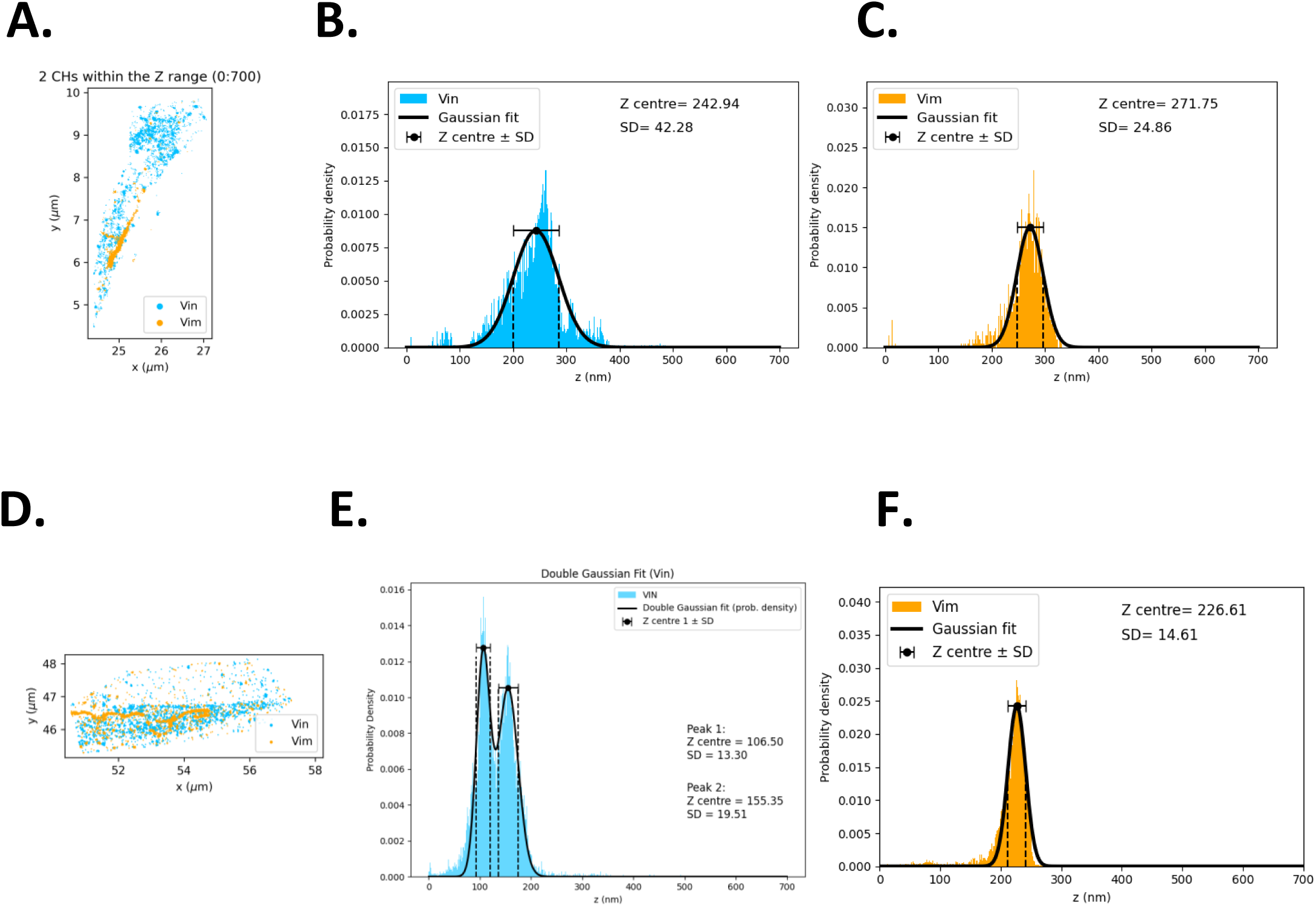
Vimentin spans multiple protein layers within the focal adhesion nanoarchitecture. (A) iPALM scatter plot of a focal adhesion showing vinculin (Vin, blue) and vimentin (Vim, orange). (B-C) Single-Gaussian fits of Z-center distributions for vinculin (B) and vimentin (C) from the adhesion in (A). (D) iPALM scatter plot of a second focal adhesion exhibiting distinct stratification. (E-F) Probability density plots for the adhesion in (D). Vinculin displays a bimodal, double-Gaussian distribution (E), while vimentin fits a single Gaussian peak (F) located above the primary vinculin layer.

**Figure S9.**
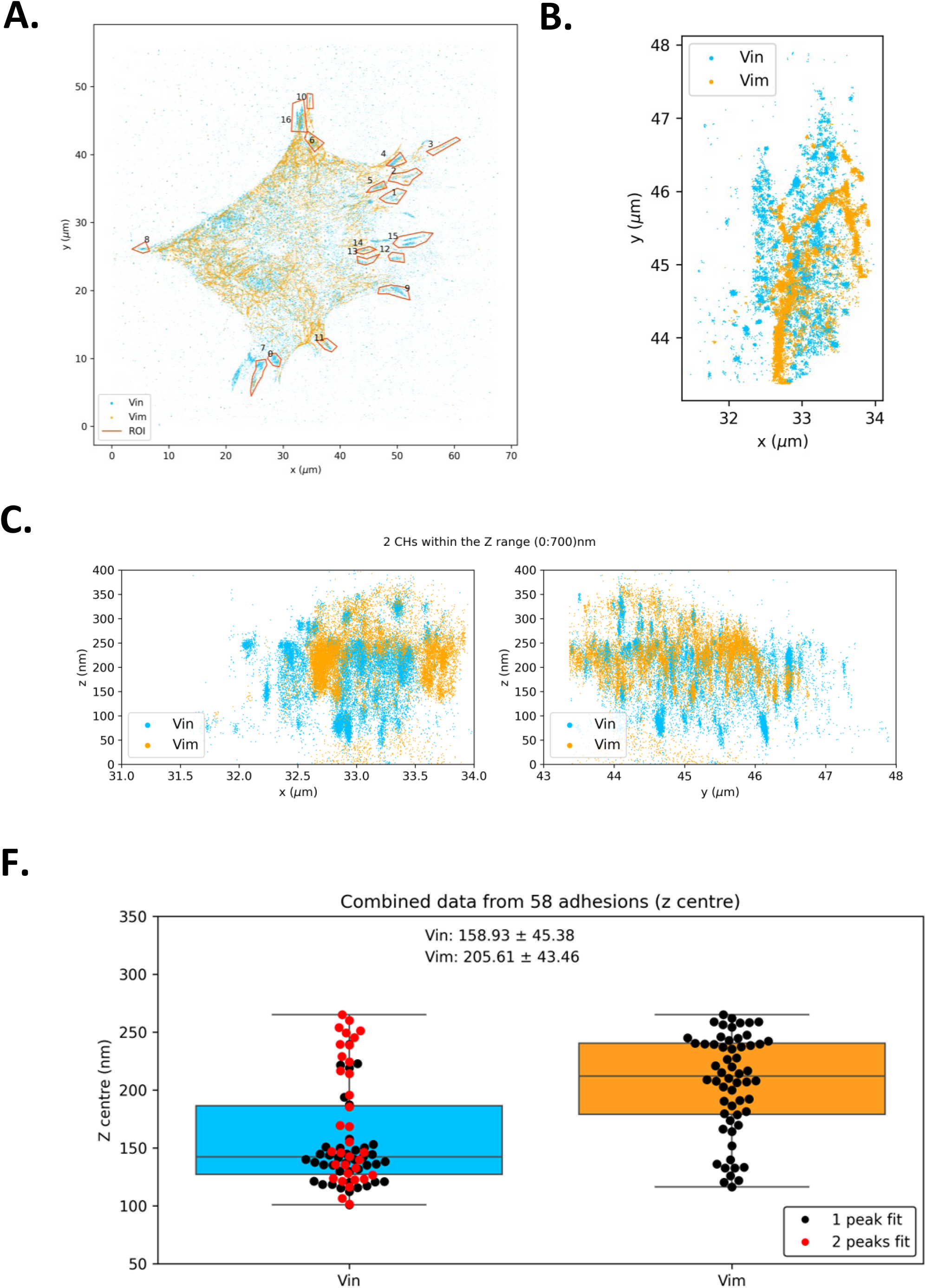
Cell-to-cell variability in vimentin localization across focal adhesion layers. (A–C) Z-center distributions of vimentin and vinculin from individual cells analyzed independently. (D) Summary plot illustrating intercellular consistency in vimentin positioning across FA layers, supporting a conserved nanoarchitectural role.

**Figure S10.**
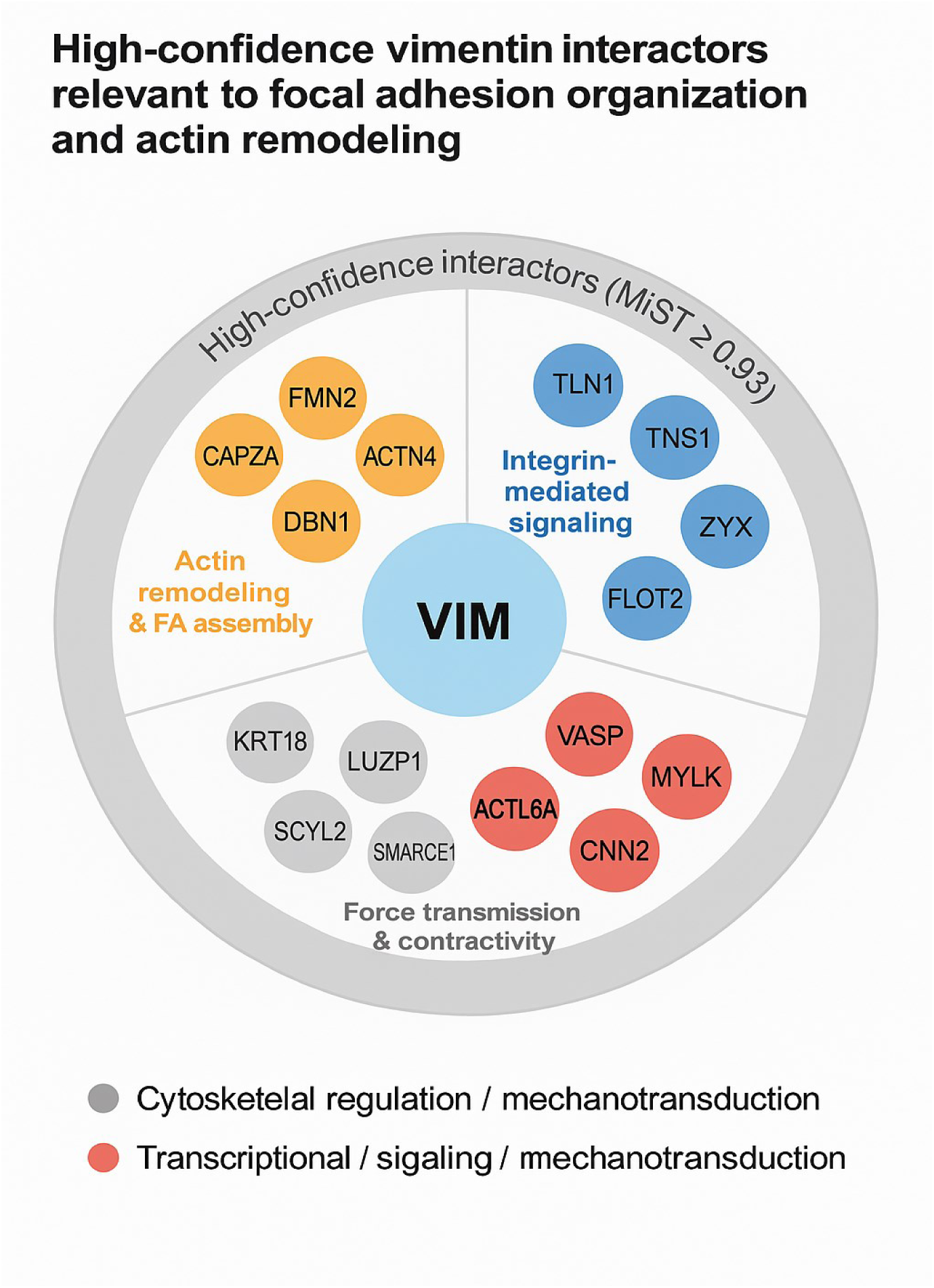
Vimentin interaction network linked to focal adhesion organization and actin remodeling. Network diagram summarizing high-confidence vimentin interactors identified by proteomics and functional annotation. Interactors are grouped by functional modules, including actin remodeling, integrin-mediated signaling, force transmission, and mechanotransduction, highlighting pathways through which vimentin regulates focal adhesion stability and organization.

